# Quantifying Treatment Resistance in Mixtures of Gastrointestinal Stromal Tumor Cells with BARMIX

**DOI:** 10.64898/2026.03.23.713602

**Authors:** Mohammad Darbalaei, Thomas Mühlenberg, Julia Zummack, Philip Dujardin, Susanne Grunewald, Anna Baginska, Patricia Munteanu, Marina Martinez Cruz, Madeleine Dorsch, Alexander Schramm, Sebastian Bauer, Daniel Hoffmann, Barbara M. Grüner

## Abstract

Targeted therapies in gastrointestinal stromal tumors (GIST) often fail due to heterogeneous resistance mutations arising across metastatic sites. Efficient, rational design of mutation-specific therapies requires the ability to quantify treatment resistance across many genotypes in parallel. Here, we present BARcode MIXture analysis (BARMIX), a platform combining multiplexed experiments with DNA-barcoded cancer cell mixtures *in vitro* and *in vivo*, and a probabilistic framework for quantitative assessment of genotype-specific treatment resistance. BARMIX efficiently and accurately recapitulated known clinical resistance patterns in GIST and matched resistance measurements from individual cell lines *in vitro* and *in vivo*. This experimental-computational approach provides a scalable and broadly applicable strategy for quantifying treatment responses in complex cell populations, enabling systematic preclinical testing of new drugs and combinations to identify mutation-specific therapeutic options for precision oncology in GIST and beyond.

## INTRODUCTION

Gastrointestinal stromal tumors (GIST) are in ~80% of cases driven by activating mutations of the KIT receptor tyrosine kinase and targeted therapies against mutant KIT have revolutionized the treatment, significantly improving patient outcomes^1–4^. Imatinib, a selective KIT inhibitor, approved as first-line therapy in KIT-mutant GIST, effectively inhibits oncogenic signaling in most tumors harboring primary mutations, most commonly those located in the juxtamembrane domain^1^. However, despite initial clinical responses, most patients eventually develop imatinib resistance, typically within two years of treatment initiation^5^. Secondary resistance to imatinib is mostly conferred by secondary mutations within the KIT kinase domain, leading to treatment failure and disease progression^5^. A subgroup of KIT/PDGFRA mutant GIST also harbor mutations that activate KIT-dependent signaling pathways irrespective of upstream activation through KIT (e.g. NF1, PI3K, BRAF, KRAS)^6, 7^. We and others have previously shown that oncogenic variants of these KIT-downstream signaling intermediates have been found in samples of patients with tyrosine kinase inhibitor (TKI) resistance and that these variants confer resistance when modeled *in vitro*^8^.

Overcoming resistance is further complicated by both inter- and intra-patient heterogeneity – as no current drug is able to inhibit the full spectrum of resistance mutations^8–11^. Both, the resistance profile of individual patients, the typical profile of the general cohort but then also the specific inhibitory spectrum of novel drugs are needed to develop effective salvage subsequent lines of treatment. Preclinical research has long been hampered by a scarcity of primary cell lines representative of this heterogeneity. We therefore recently developed an isogenic GIST cell line panel based on the GIST-T1 cell line with a primary KIT activation mutation. This panel comprises resistance mutations which recapitulates most of the genomic heterogeneity of resistance that is mediated through KIT or its oncogenic signaling pathways^8, 12, 13^. By introducing clinically relevant mutations into a common genetic background, these models enable controlled interrogation of how specific alterations confer differential sensitivity or resistance to targeted therapies. This allows direct comparison across diverse resistant genotypes, reflecting the mutational diversity found in refractory GIST patients, while minimizing confounding factors arising from variable genetic backgrounds.

However, current preclinical *in vivo* modeling of treatment resistance in GIST relies heavily on cell line derived xenograft or patient-derived xenograft (PDX) models, which, although valuable, are resource-intensive, low throughput, and insufficiently quantitative when evaluating heterogeneous tumor cell populations^14–17^. Advances in DNA barcoding technology have enabled multiplexed tracking of distinct tumor clones within pooled cell populations, permitting simultaneous interrogation of treatment responses across multiple genotypes^18–20^. *In vivo* lineage tracing via DNA barcoding permits high-resolution quantification of clonal dynamics, including baseline fitness differences, genotype-dependent growth behavior, and responses to therapeutic pressure, yet its application in modeling resistance has been limited, often lacking integration with robust statistical frameworks for quantitative resistance assessment^19–21^.

In fact, several studies have used DNA barcoding to measure clonal responses to therapy by tracking temporal changes in relative barcode frequencies^20, 22–24^. Although these approaches have provided valuable insights into competition and resistance mechanisms, they rely solely on relative abundances, which are suitable to reflect compositional changes^25, 26^ but in general not absolute clone-specific growth or treatment response. Because all relative frequencies sum to one, an increase in one clone’s proportion necessarily coincides with a decrease in others, even if all clones are growing. Conversely, a clone that expands more slowly than its competitors may appear to shrink despite increasing in absolute cell number. This coupling between clones prevents direct interpretation of relative abundance changes in terms of true growth behavior or drug response.

To overcome these limitations, we developed a comprehensive high-throughput platform that combines multiplexed DNA-barcoded GIST cell line pools with a new Bayesian modeling framework that integrates barcode and volume data to quantify treatment resistance at the clonal level. We established a scalable experimental platform and benchmarked it against known clinical resistance patterns, and we further validated it by single-cell line experiments. Using this integrated methodology, we directly interrogated eight isogenic GIST cell lines harboring diverse secondary resistance mutations, both within KIT and across downstream signaling pathways, and tested it in technical triplicates against multiple targeted therapies *in vitro* and *in vivo*. This approach not only recapitulated established response and resistance profiles to first-line imatinib, as well as second- and third-line agents and combination treatments but also allowed us to model whether cellular behavior is maintained in mixed populations compared to individual cultures. By reducing experimental scales while enhancing quantitative power and biological resolution, our platform advances current preclinical modeling paradigms and provides a critical tool to inform precision oncology strategies in GIST and other malignancies characterized by mutational heterogeneity and acquired resistance.

## RESULTS

### Quantitative treatment resistance in barcoded cell line pools reflects individual cell line responses

To systematically dissect genotype-specific drug resistance in GIST, we established a scalable high-throughput platform that is based on barcode-labeled cell line variants for pooled therapeutic profiling. First, we used retroviral expression to stably insert unique 6-nucleotide DNA barcodes^27^ (Extended Data Fig. 1a) into the patient-derived GIST cell line GIST-T1 and each of seven CRISPR-edited sublines bearing additional mutations either within KIT (T1-V654A, T1-D816A, T1-D816E, T1-A829P) or its downstream effectors (T1-TSC2, T1-PTEN, T1-G12R-HOM). Barcode-based tracking by high-throughput sequencing (HTseq), following pooled drug exposure *in vitro* or *in vivo*, enabled computational deconvolution of genotype-specific responses and clonal dynamics. For each genotype, we generated three cell lines, each labeled with a different barcode, and in this way realized three internal replicates per genotype (Fig. 1a; Extended Data Fig. 1b) to account for sources of variability in individual experiments.

**Fig. 1:**
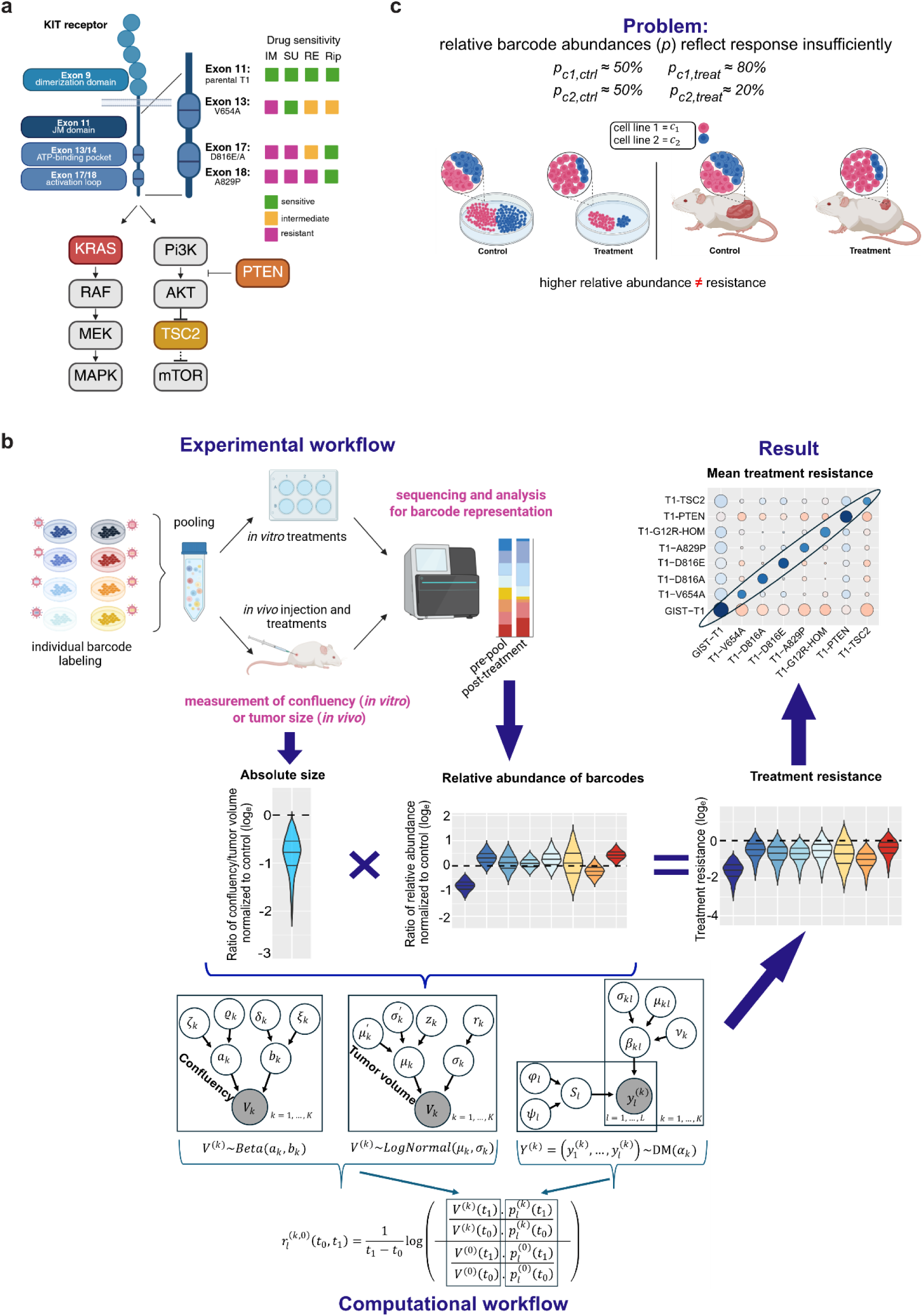
Combination of an isogenic cell line model with molecular barcoding to analyze genotype-specific drug resistance mutations in GIST. **a)** Schematic overview of the KIT receptor and secondary resistance mutations in KIT or its downstream signaling intermediates as represented in the cell line pools. For secondary mutations in KIT response patterns to approved KIT-inhibitors imatinib (IM), sunitinib (SU), regorafenib (RE), and ripretinib (RI) is shown. **b)** Schematic of experimental and computational workflows. Hierarchical joint Bayesian inference of drug resistance (result) combines measured tumor volumes (*in vivo*)/confluencies (*in vitro*) and relative barcode abundances (middle panel). **c)** Schematic examples of insufficiency of relative abundances. Left: both cell lines (c1, red; c2, blue) decrease in confluency under treatment, i.e. both are sensitive. However, relative abundances are diverging (increase of c1, decrease of c2). Right: tumor shrinks under treatment, i.e. both cell lines are sensitive to treatment, but there is no effect on relative abundances.

To assess genotype-specific responses in direct comparison, all eight cell lines with their three independent technical replicates can be pooled and exposed to a treatment *in vitro* or *in vivo.* Afterwards, samples are sequenced for barcode representation to assess each genotype’s response to the treatment (Fig. 1b, upper panel). However, relative barcode abundances assessed by HTseq of pooled barcoded cell lines are “count-compositional”^26^ and can only measure relative treatment effects in a pool, not absolute treatment effects on individual cell lines. To infer the latter, we must include volumetric parameters of the total pool size, namely confluency (*in vitro*) or tumor volume (*in vivo*) into the equation (Fig. 1c). The framework follows a probabilistic paradigm and propagates uncertainties of the different measurements through hierarchical Bayesian models to a joint posterior probability distribution of the quantitative treatment resistance (QTR) for each clone-drug pair (Fig. 1b, lower panel).

One key assumption of our model is that barcoding does not alter cell-intrinsic growth or treatment responses, and that barcoded lines behave comparably in pooled and individual settings. To validate this, we first compared morphology and relative growth of barcoded versus non-barcoded cell lines, with and without imatinib, using experimentally derived parameters in a computational model (Fig. 2a). As imatinib is the first-line therapy for GIST^13^, GIST-T1 is expected to respond at 100 nM, whereas lines carrying secondary resistance mutations should remain insensitive. Consistent with this, barcoding did not introduce differences beyond normal biological variation in morphology, growth, or imatinib response (Fig. 2b, Extended Data Fig. 2a).

**Fig. 2:**
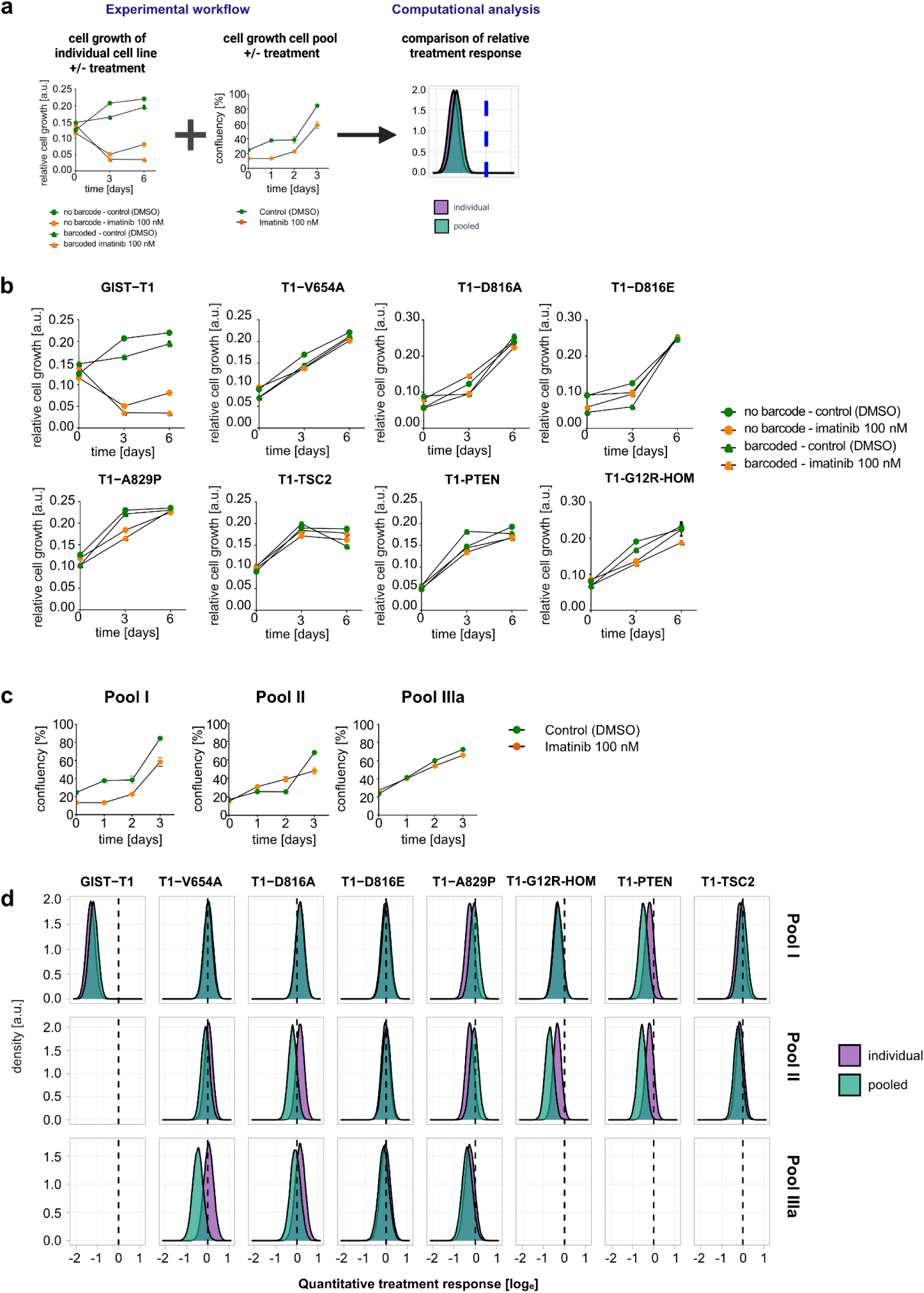
Quantitative assessment of cell growth of barcoded versus non-barcoded cells with and without treatment. **a)** Relative cell growth of individual cell lines as well as cell pools with and without treatment was measured experimentally with viability assays and integrated for the analysis of quantitative treatment response of each genotype alone (individual) or in a cell line mix (pooled). This allows the assessment of potential impact of cell lines/genotypes on each other. **b)** Relative growth of individual cell lines without and with barcode labels under control or imatinib (100 nM) treatment over time. **c)** Relative growth of barcoded cell line pools under control or imatinib (100nM) treatment over time as measured by relative confluency. **d)** Density plots of the posterior distributions of treatment responses (natural-log scale). Columns represent genotypes, rows represent pools. Purple and green distributions represent results from individual and pooled experiments, respectively. The vertical black dashed line marks the threshold separating treatment-sensitivity (left) and resistance (right).

We next assessed whether individual behavior was preserved in competitive contexts by generating three barcoded pools: pool I (all eight cell lines, each in three barcode variants), pool II (all lines except parental GIST-T1), and pool IIIa (only secondary KIT-mutant lines: T1-V654A, T1-D816A, T1-D816E, T1-A829P). Each pool was treated for three days with vehicle or imatinib. As expected, confluency changes were minimal, as only a small fraction of cells, such as GIST-T1 in pool I, were imatinib-sensitive (Fig. 2c).

Barcode abundance was quantified by HT-seq at seeding (“pre-pool”) and after three days of treatment. From these measurements, we inferred probability distributions of quantitative treatment resistance (QTR) to 100 nM imatinib for each cell line in each pool (Fig. 1c; Extended Data Fig. 2b). In parallel, individual cell-line assays were analyzed using the volumetric branch of the computational framework, yielding corresponding QTR distributions. This enabled direct comparison of QTRs derived from pooled versus individual assays (Fig. 2d). For all but one cell line, pooled and individual QTR distributions showed quantitative agreement, defined as a shift smaller than the uncertainty of either distribution and consistent directionality (sensitivity vs resistance). Minor deviations were observed for some genotypes, particularly in pool II, but only T1-V654A in pool IIIa violated quantitative agreement: pool IIIa suggested a negative QTR (imatinib sensitivity), whereas both the individual assays and pools I and II indicated no response, with posterior QTR centered near zero. Although rare, such inconsistencies underscore the need for benchmarking, as they may otherwise lead to erroneous interpretations. The isolated discrepancy in pool IIIa may reflect a technical issue specific to that pool or a context-dependent interaction among KIT-mutant lines.

### Barcoded pools enable multiplexed drug testing and ranking

To scale the platform for high-throughput application, we assembled three pooled mixtures: pool I containing all genotypes, pool II excluding the imatinib-responsive parental GIST-T1 line, and pool IIIa comprising only secondary KIT-mutant lines. Barcode representation was quantified at seeding and after three days of treatment by next-generation sequencing, and confluency was measured at both time points. These data served as inputs to our Bayesian framework to infer quantitative treatment responses (Fig. 3a).

**Fig. 3:**
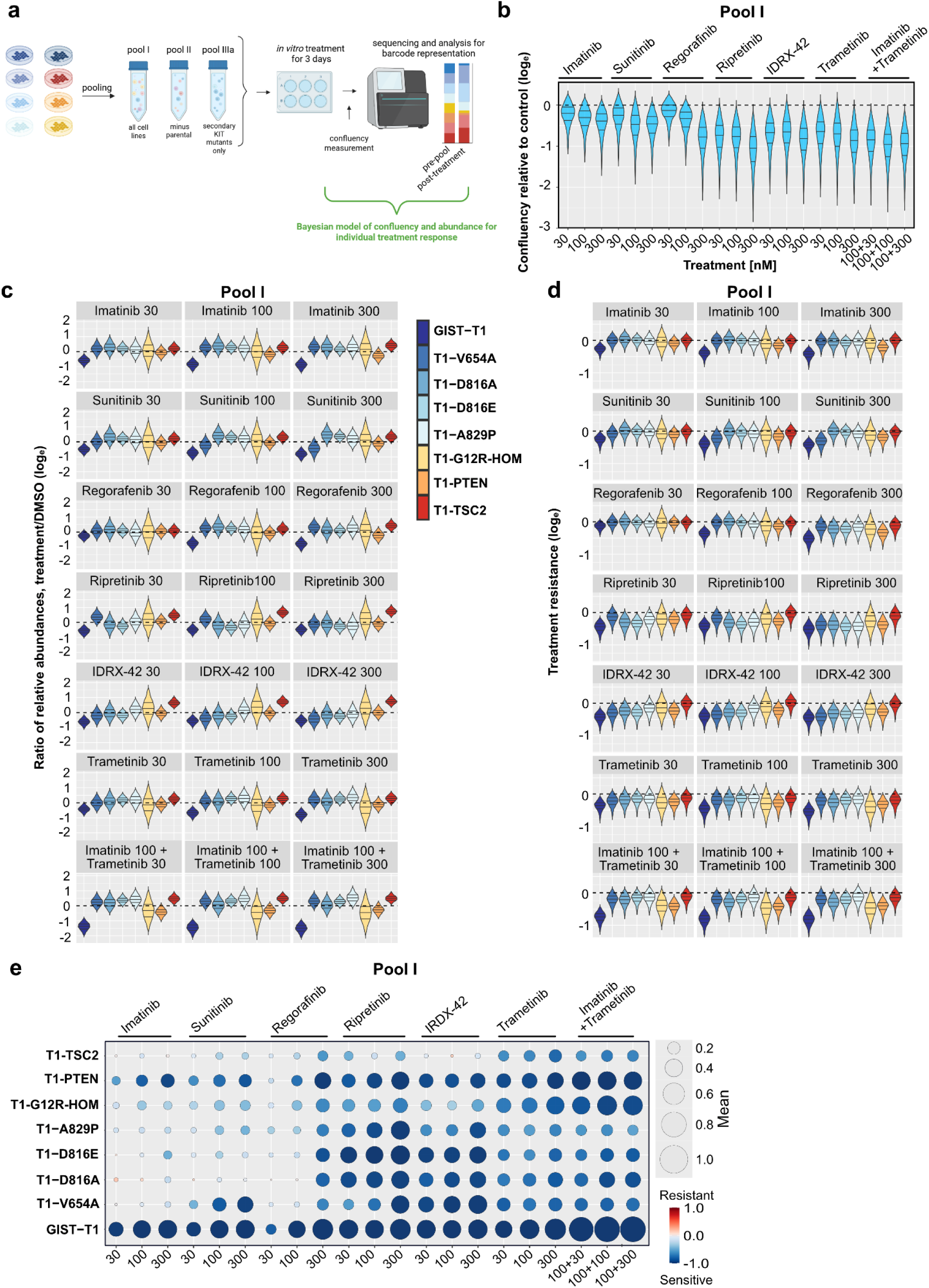
Quantitative assessment of short-term treatment responses *in vitro*. **a)** Barcoded cell lines were pooled and treated with different compounds, with confluency measured and barcode sequencing performed at 0 and 72 hours. Pools I (all cell lines), II (all cell lines except GIST-T1), and IIIa (only secondary cKIT resistance mutations T1-V654A, T1-D816A, T1-816E, and T1-A829P) were assembled as indicated. **b)** Natural-log of ratio of confluency between treated and control conditions in pool I. Uncertainty is inferred from our Bayesian model of confluency measurements (Fig. 1c and Methods). Dashed line marks baseline (log ratio = 0), corresponding to equal confluency under treatment and control. **c)** Ratio of relative abundance for each genotype within Pool I between treated and control conditions. Uncertainty is inferred from our Bayesian model of relative barcode abundance (Fig 1c and Methods). **d)** Quantitative Treatment Resistance (QTR) for Pool I is inferred by integrating Bayesian models of cell confluency and relative barcode abundance, with uncertainty propagated from both components. Each distribution reflects resistance on a natural-log scale for a given genotype–treatment pair. Dashed line separates sensitive (<0) and resistant (>0) ranges of QTR. **e)** Bubble plot summarizes treatment resistance across all cell lines in Pool I, facilitating direct comparison of all parameters. Bubble sizes reflect strengths of QTR (posterior mean), while color represents the direction of QTR: blue indicates sensitivity, i.e. treated cells growing slower than in control; red indicates resistance, i.e. treated cells growing faster in control.

Using pool I, we treated cells with imatinib, sunitinib, regorafenib, ripretinib, IDRX-42, trametinib, and imatinib-trametinib combinations across 30-1000 nM. These compounds span all approved treatment lines for KIT-mutant GIST, as well as agents currently in late-stage clinical evaluation. IDRX-42 is in phase 3 trials (NCT07218926), and the sunitinib + bezuclastinib combination is under phase 3 investigation (PEAK, NCT05208047). Trametinib, a MEK inhibitor with known toxicity in GIST, was included as a tool compound to probe downstream pathway inhibition. Confluency was measured at treatment start and after three days (Extended Data Fig. 3a), and relative confluency - including uncertainty - was estimated using a Bayesian model (Fig. 3b). In parallel, barcode abundances were quantified in pre-pools and post-treatment pools by sequencing (Fig. 3c). Integrating both data types in our joint model yielded QTR estimates, with uncertainties, for each cell line across all treatments (Fig. 3d).

To enable direct comparison across genotypes and drug conditions, we visualized QTR values using bubble plots, where bubble size reflects mean QTR and color encodes the credibility of the effect direction (dark blue to dark red) (Fig. 3e; Extended Data Fig. 3b). The same analysis was applied to pools II (Extended Data Fig. 3c–g) and IIIa (Extended Data Fig. 3h–k). For pool II, we focused on imatinib, sunitinib, ripretinib, regorafenib, and IDRX-42 to cover first- to fourth-line therapies and a late-stage investigational agent. For pool IIIa, containing only secondary KIT-mutant genotypes, we analyzed imatinib and the most recent KIT-targeted inhibitors (ripretinib, regorafenib, IDRX-42). These additional pools allowed us to test whether the presence of the imatinib-responsive parental GIST-T1 line influences drug effects on other genotypes, and whether BARMIX can resolve subtle differences among closely related KIT-mutant variants.

Across pools, QTR estimates for each cell line–treatment pair were highly consistent, demonstrating reproducibility and indicating that BARMIX performance is robust to pool composition. As expected, GIST-T1 was the most sensitive line across all inhibitors (Fig. 3e). All secondary KIT-mutant lines were markedly resistant to imatinib, with T1-PTEN showing residual sensitivity as previously reported ^8^. Sunitinib selectively inhibited the ATP-binding pocket mutant T1-V654A, whereas activation-loop mutants (T1-D816A, T1-D816E, T1-A829P) remained resistant. Conversely, ripretinib strongly suppressed activation-loop mutants, while T1-V654A showed reduced sensitivity across pools. Lines with downstream KIT-pathway mutations showed minimal response to KIT inhibitors in pools I and II but were sensitive to MEK inhibition by trametinib, with enhanced effects in combination with imatinib. Notably, IDRX-42 exhibited strong activity across all secondary KIT-mutant lines and showed partial activity in downstream-mutant lines, supporting its potential as a treatment option in multi-resistant GIST.

As a simplified and more direct comparison of drug effect on genotypes that allows BARMIX to easily identify best compound-genotype combinations, we implemented a ranking system that allows to easily identify the best performing drug and concentration for each genotype. This system ranks treatments based on the posterior predictive median of treatment resistance for concentrations in a range that seems clinically achievable (Fig. 4a-h; for a complete ranking see Extended data Fig. 4a). Again, ripretinib at 30 nM was most efficient for the three cell lines harboring secondary AL mutations, whereas AP-mutant T1-V654A showed a higher QTR but was most sensitive to IDRX-42 at 100 nM and even 30 nM. Interestingly, even though GIST-T1 was sensitive to imatinib at 100 nM, a combination with trametinib at 30 nM proved to be the most efficient for this cell line, and this also was true for the three downstream-signaling mutant cell lines (T1-PTEN, T1-TSC2, and T1-G12R-HOM). We applied this ranking system to pools II and IIIa (Extended data Figs. 4b-e) which were treated with selected drugs as described above. Here, IDRX-42 showed the best performance for all imatinib resistant cell lines, except T1-TSC2 which was highly resistant against direct KIT inhibition. While we observed minor quantitative shifts between ranking tables between pools, the main conclusions did not depend on the analyzed pool.

**Fig. 4:**
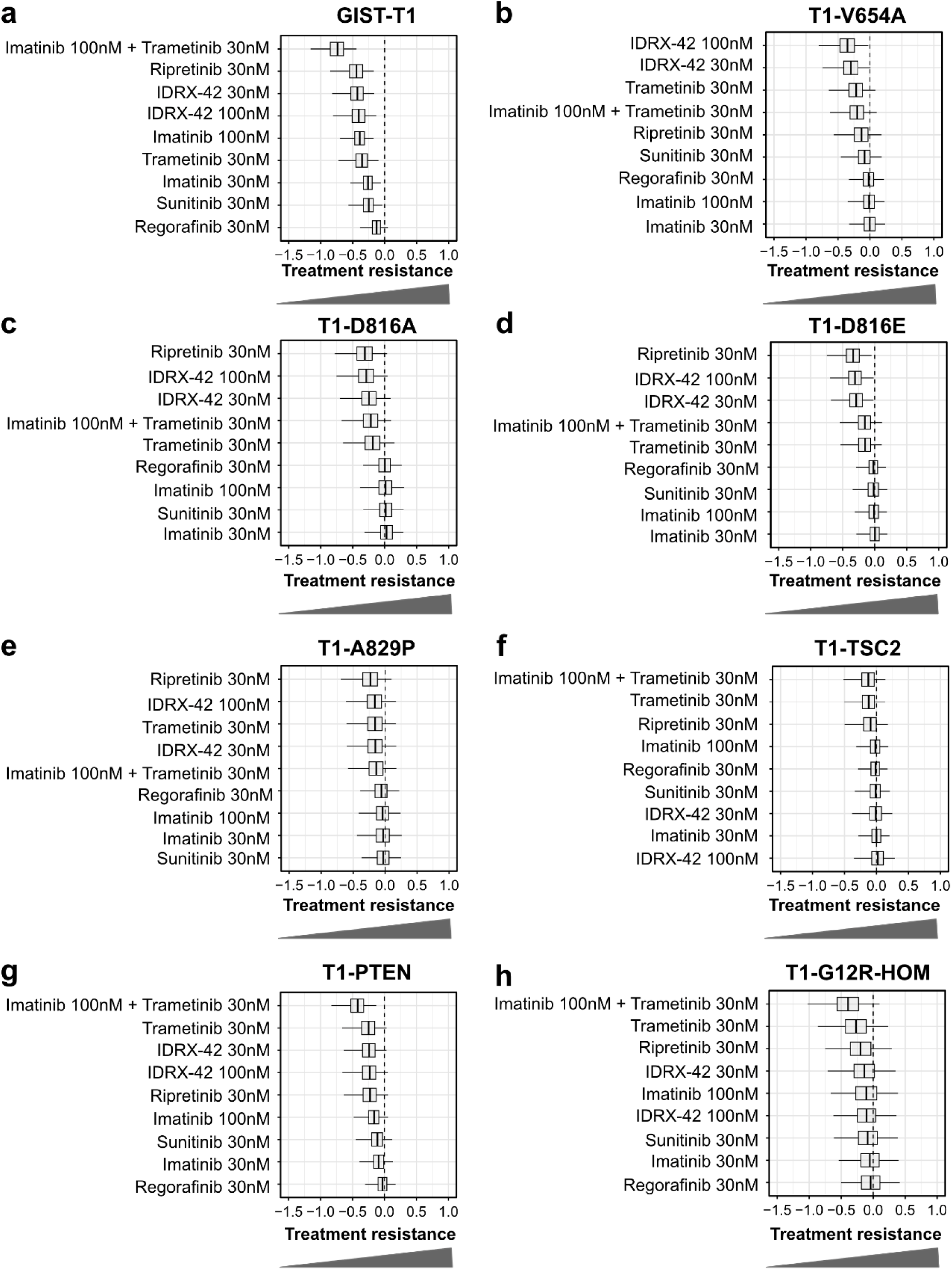
Quantitative ranking system to support preclinical decision making. **a-h)** Vertical axes: selected drugs with their respective concentrations representing clinically relevant doses. Drugs are ranked for each genotype in Pool I as indicated by the posterior predictive median of quantitative treatment resistance (natural-log scale, horizontal axes). The most favorable drug treatment (highest sensitivity) for each genotype appears on top. Box plots summarize the posterior predictive distribution: boxes represent the interquartile range (IQR), horizontal lines mark the median, and whiskers extend to the 10th and 90th percentiles. The vertical dashed line at zero separates treatment-sensitivity (left) and resistance (right).

We then sought to analyze whether our platform could be applied to more complex long-term treatment settings. To this end, we seeded pools I, II, and IIIb, harvested a fraction of the cells after 7 days for analysis and reseeded similar cell numbers again for another 7 days of treatment. Notably, this time pool III additionally contained the GIST-T1 cell line (pool IIIb), to serve as an internal positive control for treatment response. For treatment start at day 0, day 7, and day 14 confluency was measured, and relative barcode abundances were analyzed (Fig. 5a). These data were combined in our computational model for all three pools as described for the short-term experiment and QTR was assessed for all treatments at both time-points (Figs. 5b-e and Extended data Figs. 5a-di).

**Fig. 5:**
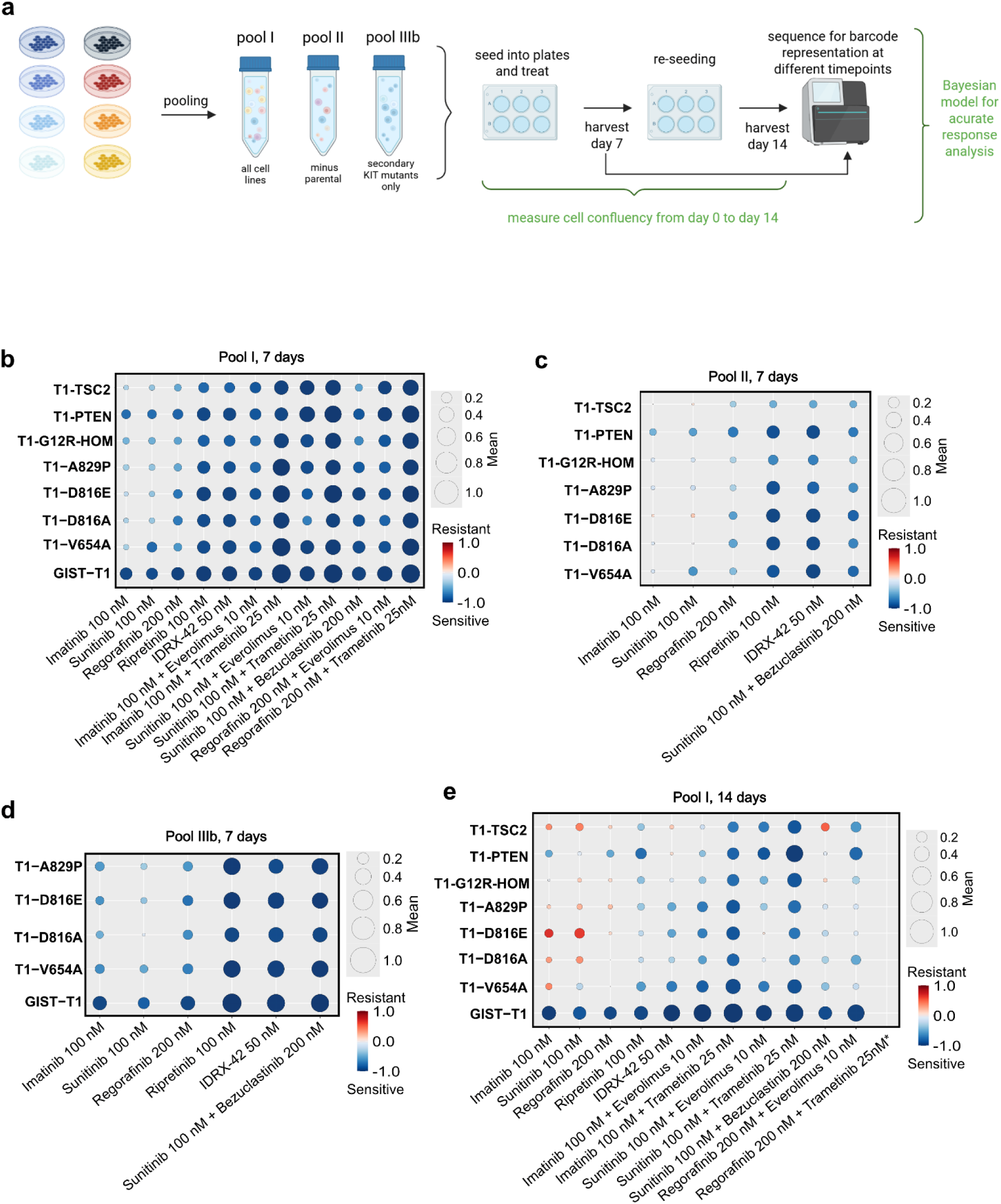
Quantitative assessment of *in vitro* long-term treatment responses. **a)** Schematic of the *in vitro* workflow. Barcoded cell lines were pooled as indicated and treated for 14 days. After 7 days, cells were harvested and 100,000 cells were reseeded for another 7 days of treatment. Confluency measurements and barcode sequencing were performed for all pools at days 0, 7 and 14. **b-e)** Results are shown for Pool I **(b, e),** Pool II **(c),** and Pool IIIb **(d)** at day 7 **(b-d)** and day 14 **(e)**. Bubble plots summarize quantitative treatment resistance (QTR) across all cell lines within each pool, enabling direct comparison across treatments and concentrations. Bubble size represents the posterior mean QTR, and color indicates the direction of effect: **blue denotes resistance** (treated cells grow faster than in control), whereas **red denotes sensitivity** (treated cells grow slower than in control).

In pool I, GIST-T1 exhibited the highest sensitivity to all treatments at both day 7 and 14. The T1-V654A variant, while resistant to imatinib, remained sensitive to sunitinib. In contrast, T1-PTEN consistently displayed the lowest QTR among all imatinib-resistant cell lines. Comparison across time points revealed that the 7-day results were largely concordant with those obtained at 3 days, albeit more pronounced. At 14 days, treatment effects were further amplified, although in several instances positive QTR values were observed, indicating that certain cell lines proliferated more efficiently under treatment than under control conditions. Notable exceptions to the general trend included unexpectedly low QTR values for T1-TSC2 and T1-G12R in response to ripretinib (100 nM), as well as an unanticipated QTR for T1-PTEN upon exposure to IDRX-42. However, given the potential bias introduced by splitting and re-seeding, no further 14-day assays were pursued. In pool II, comprised of the seven imatinib-resistant cell lines, all lines displayed strong QTR to imatinib, as expected. As before, T1-V654A showed resistance to sunitinib under the same treatment in pool II. In pool IIIb, which included only cell lines harboring primary and secondary KIT mutations, T1-V654A again showed higher imatinib resistance than anticipated, which might be explained by our previous findings that this cell line in this pool is more resistant, when in direct comparison to only KIT secondary mutants (Fig. 2d). Furthermore, in line with previous findings, all cell lines remained sensitive to ripretinib, IDRX-42, and the combination of bezuclastinib with sunitinib.

### BARMIX enables quantitative preclinical modelling in vivo

To extend BARMIX to *in vivo* settings, we transplanted pools I, II, and IIIb into mice and, after tumor establishment, treated animals with selected compounds. Imatinib and sunitinib served as benchmark inhibitors to validate the *in vivo* model, and in pool I we additionally tested trametinib alone or in combination to probe KIT-downstream inhibition in genotypes with reduced KIT dependence (T1-G12R-Hom, T1-PTEN, T1-TSC2). Tumor volumes were recorded, and tumors were harvested for barcode sequencing alongside the pre-injection pools (Fig. 6a). For pool I, five treatments plus vehicle control were tested (Fig. 6b). Because changes in barcode abundance may reflect either treatment effects or intrinsic *in vivo* growth differences, each experimental setup included an additional mouse sacrificed at treatment onset to establish baseline representation. We quantified endpoint tumor volumes (Fig. 6c) and calculated treatment-to-control volume ratios (Fig. 6d), which replace the *in vitro* confluency ratios in our computational framework. These ratios were integrated with barcode abundances in a joint Bayesian model (Extended Data Fig. 6a) to infer QTR values for each genotype and treatment *in vivo* (Extended Data Fig. 6b). Results were visualized using bubble plots for direct comparison across all genotype-drug combinations (Fig. 6e; Extended Data Fig. 6c), and treatment ranking per genotype facilitated identification of the most effective therapy (Extended Data Fig. 6d).

**Fig. 6:**
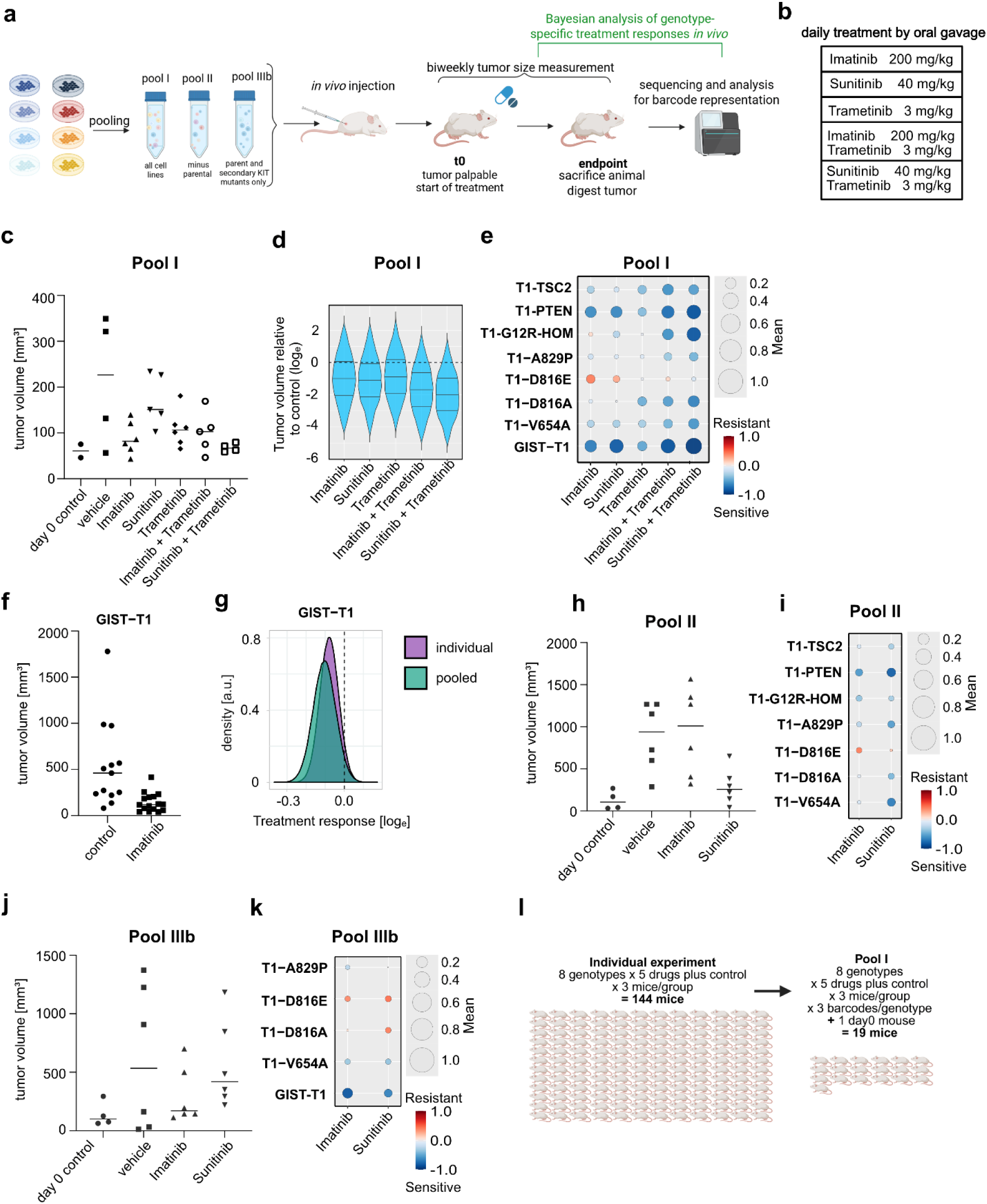
BARMIX enables quantitative preclinical modelling *in vivo*. **a)** Schematic of *in vivo* workflow. Barcoded cell lines were pooled as indicated and injected into mice. Tumor volume was measured at treatment start (*t_0_*) and endpoint, and barcode sequencing performed at *t_0_* and endpoint. **b)** Drugs were administered daily by oral gavage. Single and combination treatments are listed with corresponding dosages. **c)** Tumor volume measurements in Pool I for the indicated treatment conditions and time points. **d)** Natural-log scaled ratio of tumor volumes, treated / control, in pool I. **e)** Summary of *in vivo* quantitative treatment resistance (QTR) across genotypes Pool I. Bubble size reflects the mean QTR, color shading represents the direction: blue indicates sensitivity (treated cells grow slower than in control); red indicates resistance (treated cells grow faster than in control). **f)** Tumor volumes in individual GIST–T1 xenografts after 14 days of Imatinib treatment. **g)** Density plots of the posterior distributions of treatment response (natural-log scale) for GIST–T1. Purple and green distributions represent results from individual and pooled experiments, respectively. The vertical black dashed line separates treatment-sensitive (left) and resistant (right). **h, j**) Tumor volume measurements in Pool II **(i)** and Pool IIIb **(k)** across treatment groups. **i, k**) Summary of treatment resistance across genotypes and drug conditions in Pool II **(i)** and Pool IIIb **(k)**. **l)** Comparison of animal numbers required for conventional individual experiments (left) and barcoded pool design (right).

As expected, in pool I, which contained all investigated cell lines, GIST-T1 responded to all treatments (lowest QTR). While this cell line was least sensitive (highest QTR) to trametinib alone, the combination of trametinib and sunitinib decreased QTR most potently. In concordance with the *in vitro* results, cell lines with secondary KIT mutations, as well as those with downstream mutations displayed treatment resistance to imatinib and sunitinib alone, which was attenuated in combinations with trametinib. Only T1-PTEN was inhibited by imatinib and sunitinib alone, indicating a sustained dependency of this genotype on KIT signaling.

To confirm that the platform yields robust results while significantly downscaling *in vivo* experiments and increasing throughput, we benchmarked pooled QTR readouts against individual *in vivo* treatments of GIST-T1 tumors exposed to imatinib (Fig. 6f). QTR values from single-line experiments were highly congruent with those from barcoded and pooled settings, demonstrating that neither barcoding nor pooling altered *in vivo* growth behavior or treatment response (Fig. 6g). We next assessed whether pool composition influenced outcomes by applying the platform to two additional pools with overlapping cell-line compositions, pool II (Fig. 6h,i; Extended Data Fig. 6e–i) and pool III (Fig. 6j,k; Extended Data Fig. 6k–o).

Mice injected with pools II and IIIb were treated with imatinib, sunitinib, or control. In pool II, which included all lines except GIST-T1, all barcoded lines showed strong resistance to imatinib, with T1-PTEN displaying the lowest QTR under both treatments. Among secondary KIT mutants, T1-V654A remained slightly more responsive to sunitinib and showed minimal residual sensitivity to imatinib. Pool IIIb, composed exclusively of KIT-mutant lines plus parental GIST-T1, reproduced the expected patterns: all secondary KIT mutants were resistant to both drugs, whereas GIST-T1 remained imatinib-sensitive. Across all three *in vivo* pools, BARMIX consistently delivered quantitative, reproducible treatment-response profiles in a multiplexed format. Notably, pool I required only 19 mice (three per treatment arm plus one baseline), whereas equivalent single-line experiments would require at least 144 animals, not accounting for the larger group sizes typically needed to capture biological variability (Fig. 6l). Taken together, these results establish BARMIX as a precise, scalable, and resource-efficient approach for assessing treatment responses in complex *in vivo* settings.

## DISCUSSION

Genetic heterogeneity is a primary driver of therapeutic failure in cancer, and GIST exemplify this challenge, as resistance emerges through diverse secondary KIT mutations that confer distinct sensitivity/resistance profiles to approved TKIs^5, 13, 28^. Despite multiple approved treatment options, therapy sequence remains fixed until failure of 4^th^ line TKI ripretinib^29^. After this, selection of salvage treatments remains largely empirical since preclinical systems capable of quantitatively resolving resistance across heterogeneous tumor populations remain limited.

Modeling this diversity of resistance is essential for precision oncology and drug development. However, conventional single-genotype approaches scale poorly with genetic complexity. Here, we introduce a multiplexed barcoding strategy combined with probabilistic deconvolution that enables quantitative analysis of pooled drug responses in GIST systems *in vitro* and *in vivo*.

A central advance of this work lies in its computational framework. Our Bayesian framework models QTR as a continuous phenotype with corresponding posterior uncertainty, rather than reducing drug response to a binary sensitive or resistant classification. Bayesian inference provides a principled framework for parameter estimation and uncertainty quantification, producing posterior distributions and credible intervals that directly express the probability that a parameter lies within a given range^30^. This quantitative probabilistic output is usually more easily interpretable and less prone to misinterpretation than p-values and confidence intervals from conventional statistics^31^. Bayesian models integrate prior knowledge and, in this way, regularize against false positives. Conceptually, Bayesian modeling corresponds to learning by accumulating information, which also means that (in contrast to conventional statistics) it can and should be updated as new data become available^32^.

In our first intervention studies, we found that clinical patterns of drug sensitivity and resistance could be validated in our multiplexed model. Sunitinib and regorafenib are approved second- and third-line therapies for GIST and show activity against subsets of imatinib-resistant KIT mutations^13^. Sunitinib preferentially inhibits ATP-binding pocket (AP) mutants, whereas regorafenib is shows modest activity against several activation loop (AL) and AP mutant tumors, but in clinical practice dose levels for long-lasting therapeutic success cannot be reached^13^. Ripretinib, the first tyrosine kinase inhibitor developed specifically for KIT-mutant GIST^33^, initially demonstrated broad preclinical activity, but subsequent clinical and experimental data revealed important limitations, particularly in exon 9 and exon 11 plus 13 mutations^11, 13^. Our multiplexed *in vitro* and *in vivo* experiments generally reproduced these known clinical susceptibility patterns while simultaneously allowing interrogation of many more compounds and concentrations in direct comparison and with minimal additional effort. Doing so, our study revealed IDRX-42 as a promising option across multiple resistance mutations Although most genotype-treatment results were consistent across pools, some variation was observed, including a modest difference in the response of T1-V654A to sunitinib. Such discrepancies should be interpreted in the context of experimental scale, as animal studies must balance ethical reduction with adequate statistical power; too few animals reduce the ability to detect true biological effects and increase the likelihood that apparent differences arise from chance rather than treatment^34^. Another possible explanation is the differing pool compositions. When comparing only the KIT secondary mutation genotypes, T1-V654A consistently showed greater sensitivity than the other three cell KIT secondary mutant cell lines. However, once GIST-T1 was included in the pool, it appeared more resistant in direct comparison.

The interpretability of results with the explicit treatment of uncertainty that is delivered by BARMIX is unparalleled and easily transferable to other research questions, entities and genotypes without the need for commercial services^23^. Furthermore, *in vivo* our method reduces animal usage by several orders of magnitude, making it not only cost-and time efficient but also a more ethical option. Barcoded tumors exhibited growth behavior and drug responses similar to non-barcoded controls, supporting biological validity. Differences across pool configurations were consistent with expected sampling variability at limited cohort sizes, indicating that multiplexing preserves inferential robustness while substantially increasing experimental efficiency. At the same time, sample size remains a critical determinant of statistical power in all experimental designs, whether pooled or individual. Increasing genotypic diversity improves biological resolution but reduces effective sampling depth per clone, whereas insufficient replication increases posterior variance and thus uncertainty.

One potential limitation of BARMIX is that clone-clone interaction could distort QTR values. To detect such interactions, pool composition can be systematically varied, e.g. by selectively excluding primary or secondary resistant genotypes and then comparing resistance parameters across different pools.

Taken together, BARMIX establishes pooled tumor profiling combined with Bayesian inference as a scalable and quantitative strategy for resolving heterogeneous drug responses. The framework is extendable to RNAi-based perturbation studies, resistance evolution assays, and patient-derived models, providing a foundation for functional precision oncology in genetically complex disease.

## AUTHOR CONTRIBUTIONS

T.M., J.Z., P.D., S.G., A.B., P.M., M.M.C., Ma.D. and B.M.G. performed and analyzed *in vitro* and *in vivo* experiments. Mo.D. and D.H. designed, performed, and analyzed computational modeling. A.S. provided conceptual and experimental support and analyzed data. S.B. and B.M.G. conceptualized the project idea. Mo.D., T.M., J.Z., D.H., and B.M.G. prepared the manuscript with input from all coauthors.

## DECLARATION OF INTERESTS

T.M.; S.B.: Co-inventor of novel treatment against GIST (patent held by institution).

S.B. holds honoraria from Pharmamar, Dicephrea and Springworks, has Consulting or Advisory roles with Blueprint Medicines, Deciphera, Böhringer Ingelheim, IDRX, Adcendo, Merck, and receives funding from IRDX and von Pfeffel Pharmaceuticals.

All other authors declare no competing interests.

## ACKNOWLEDGEMENTS

We thank Eva Hahn, Nancy Meyer, Miriam Christoff, Alexandra Heidemann, and Yasmin Krausmüller for expert technical assistance and all members of the Grüner, Bauer, Schramm, and Hoffmann laboratories for helpful comments. We thank the central animal facility of the Medical Faculty at the University Duisburg-Essen (ZTL) for their support. Figures 1a-c, 2a, 3a, 5a, 6a,l, and Extended Data Fig. 1a created in BioRender by Grüner, BM. (2026) https://BioRender.com/. This work was supported by Deutsche Forschungsgemeinschaft (DFG, German Research Foundation) grants HO 1582/12-1, GRK 2762 (subprojects L1 and M1 to B.M.G. and D.H.) and an Emmy Noether Award from the German Research Foundation (DFG, GR4575/1 to B.M.G.). P.D. was recipient of a PhD fellowship from the Cusanuswerk.

## MATERIALS AND METHODS

### Cell lines and culture

GIST-T1 (RRID:CVCL_4976) were established from human metastatic GIST and contains a 57bp deletion in c-KIT exon 11 [47]. T1-A829P and T1-D816E were generated as described previously^35, 36^. GIST-T1 sublines (T1-V654A; T1-D816A; T1-PTEN; T-KRAS-G12R; T1-TSC2) were generated by CRISPR/Cas9-mediated gene editing in GIST-T1, as described previously^35^. GIST-T1 and all its sublines were grown in Iscove’s Modified Dulbecco’s Medium (IMDM) supplemented with 10% FBS (BRAND), 2 mM L-Glutamin and 50 U/mL penicillin, 50 µg/mL streptomycin and 0.25 µg/mL amphotericin B (Gibco) at 37 °C and 5 % CO_2_. All cell lines except GIST-T1 were individually maintained under imatinib treatment. All cell lines are regularly authenticated by sequencing endogenous mutations in KIT, confirmation of KIT expression, and response to KIT inhibitor treatment. During this study, all cell lines were regularly tested for mycoplasma contamination by PCR and by MycoAlert Mycoplasma Detection Kit (Lonza).

### Cryopreservation of cell lines

All cell lines were cryopreserved in early passages in cell culture medium supplemented with 20 % FBS and 10 % DMSO. Cells were stored short-term at −80 °C and long-term at −150 °C in an ultra-low temperature freezer. Frozen cells were quickly thawed and subsequently pelleted by centrifugation at room temperature (RT) and 300 x g for 5 min. Cell pellets were resuspended in culture medium and transferred to culture dishes. After 24 h culture medium was changed and cells were passaged at least once after thawing before being used in experiments.

### Generation of barcoding plasmids

The retroviral MSCV-6NBC-GFP;PGK-Puro vectors used in this work to individually barcode different mutants of the GIST T1 cell line have been generated and described previously^27^. Each vector contains a unique and distinct six nucleotide long barcode sequence, a GFP marker gene and a puromycin resistance.

### Production of retroviral particles and viral transduction of eukaryotic cells

For the production of retroviral particles for the transduction of eukaryotic cells, HEK293T cells were seeded in 12-well plates. Individual wells were transiently transfected at a confluence of 80 % with the MSCV barcoding vectors (Extended data Fig.1a) and the packaging and envelope plasmids gag/pol-retro and VSV-G-retro as described previously^27^ using the TransIT^®^-LT1 transfection reagent (Mirus). After 24 h the culture medium was changed and the virus containing supernatant was harvested after 48 h and again after 72 h and stored at −80 °C until use. For the infection of eukaryotic cells, the virus containing supernatant was thawed at room temperature and then centrifuged for 10 min at 21,300 x g. The supernatants were then added to the wells of a 12-well plate in which the target cells were grown to approximately 70 % confluence. To increase transduction efficiency the medium was supplemented with hexadimethrine bromide (brand name: Polybrene) from Santa Cruz Biotechnology at a final concentration of 10 µg/ml. 24 h after transduction, medium was changed. Transduction efficiency was confirmed by flow cytometry for GFP after 48 hours. Afterwards, the cells were selected with puromycin (1 μg/mL) and full selection (>96 %) was again confirmed by flow cytometry for GFP.

### Flow cytometry

Flow cytometry analysis of cells was performed on a FACSCelesta™ flow cytometer (BD, Becton Dickinson) to monitor the transduction efficiency and the success of the selection. Prior to analysis, cells were harvested using trypsin, centrifuged at 300 x g for 5 min and the cell pellet was resuspended in PBS containing 5 µg/ml DAPI (to allow for the discrimination of dead cells). The FACSCelesta was equipped with a blue (488 nm), a violet (405 nm) and a green (561 nm) laser. Instrument calibration was performed daily using BD FACSDiva™ CS&T Research Beads according to the manufacturer’s instructions. BD FACSDiva™ software was used for data acquisition. Further data evaluation was performed using FlowJo® software.

### Barcode identity control

DNA sequencing was performed by Microsynth Seqlab based on the Sanger chain-termination method. Sanger sequencing was performed for validation during plasmid cloning and to confirm barcode identity and cell clone purity after transduction. Sequencing results were analyzed with the Benchling (Benchling, Inc.) and BioEdit (Ibis Bioscience) software.

### Chemicals

All compounds used for *in vitro* experiments were diluted in DMSO with the final concentration of DMSO in medium at a maximum of 1 %. DMSO only was used as vehicle control. Imatinib mesylate (LC-Labs I-5508, CID: 123596), sunitinib malate (LC-Labs S-8803, CID: 6456015), trametinib (LC-Labs T-8123, CID: 11707110), everolimus (CID: 6442177), and regorafenib (CID: 11167602) were purchased from LC-Labs, bezuclastinib (HY-145557, CID: 75593308), ripretinib (HY-112306, CID: 71584930), IDRX-42 (HY-132166, CID: 155587867) were purchased from MedChem Express.

### Cell viability assay

To analyze cell viability 1 x 10^4^ cells were seeded in triplicate in 96-well plates and cultivated for 7 days in a humidified incubator with 5 % CO_2_ at 37 °C. Cell viability was measured on day 1, day 4 and day 7 using the PrestoBlue Cell Viability Reagent (Thermo Fisher Scientific) according to manufactures instruction. The cells were incubated in the viability reagent diluted 1:10 in cell culture medium for 2 h at 37 °C and absorbance was measured in a Spark multimode microplate reader (Tecan Group) at 570 nm and 600 nm. The percentage of viable cells was determined by comparing the 570-600 nm absorbance of viable cells as measured against blank wells containing only viability reagent diluted in medium. Compounds were added to wells at various concentrations in triplicate as indicated to determine their effect on cell viability compared to vehicle (DMSO) control.

### Longitudinal monitoring of cell confluence

For longitudinal monitoring of cell confluence under drug treatment, cells were seeded in 6-well plates and cell confluence was determined optically at the indicated time points. For this, the NYONE automated cell imager and its corresponding software-based cell recognition (Synentec) were used. Before each optical measurement, the culture medium was changed to remove dead cells, and the cells were re-treated with the corresponding compound as indicated. Between measurements, the plates were incubated in a humidified incubator with 5 % CO_2_ at 37 °C.

### *In vitro* short-term drug screening

To decipher the individual fitness levels of each GIST mutant, three pools with different compositions of sublines were prepared. Pool I contained all sublines, pool II all cell lines except the parental and pool III only the secondary KIT mutants. For this, all individual barcoded cell lines were harvested and mixed at equal proportions for each pool, containing three individually barcoded versions of each cell line as indicated. Pre-pool samples were kept for barcode analysis at starting point. Then, 3×10^4^ cells were seeded for each condition in triplicate and after 24 h (d0), initial confluence was determined optically using the NYONE automated cell imager. Treatment started with four concentrations per compound as indicated (30 nM, 100 nM, 300 nM and 1000 nM) and continued for 72 hours with confluency evaluated daily.

### *In vitro* competition assay

Pools were prepared as described above. 3×10^4^ cells of the pools were seeded in triplicate and cultivated for a total of 14 days under treatment as indicated. Pre-pool samples were kept for barcode analysis at starting point. After seven days the cells were passaged, reseeded and a sample was taken to analyze the pool composition at this timepoint. After 14 days, cells were harvested and all samples analyzed by Next Generation Sequencing (NGS) as described below. Confluency was measured as described at indicated time points.

### DNA Barcode Analysis

DNA isolation from cultured cells and harvested tumors from mice was performed using the Puregene Kits (QIAGEN) or the Monarch® Spin gDNA Extraction Kit (New England Biolabs) according to the manufacturer’s instructions. DNA was rehydrated in H_2_O and a NanoDrop^TM^ (Thermo Scientific) was used to determine DNA concentration. 30 – 100% of the isolated DNA was PCR-amplified using primers that introduced the Illumina sequencing primer binding sites and adapters as well as multiplexing tags in a single PCR reaction. For the amplification the OneTaq® DNA Polymerase (New England BioLabs) and 30 PCR cycles (95 °C for 30 s, 66 °C for 30 s and 72 °C for 60 s) were chosen. PCR amplicons were separated on agarose gels and purified using the QIAquick Gel Extraction Kit (QIAGEN). PCR products were eluted twice in 10 – 30 μl ddH_2_O and concentration was measured using the Qubit^TM^ 1X dsDNA HS assay kit (Invitrogen). The molarity of each sample was determined to calculate the library concentration and pool the samples accordingly. Subsequently, the samples were pooled in equimolar concentrations and mixed with 40% PhiX control. The Qubit^TM^ 1X dsDNA HS assay kit (Invitrogen) was used to determine DNA concentration for the pooling of the libraries. Each library was sequenced using a MiniSeq Mid Output Kit on an Illumina MiniSeq sequencer. Reads per barcode per sample were extracted from the fastq files and pre-experiment to post-experiment barcode ratios were automatically calculated by custom-designed python codes^27^.

### Barcode sequencing primers

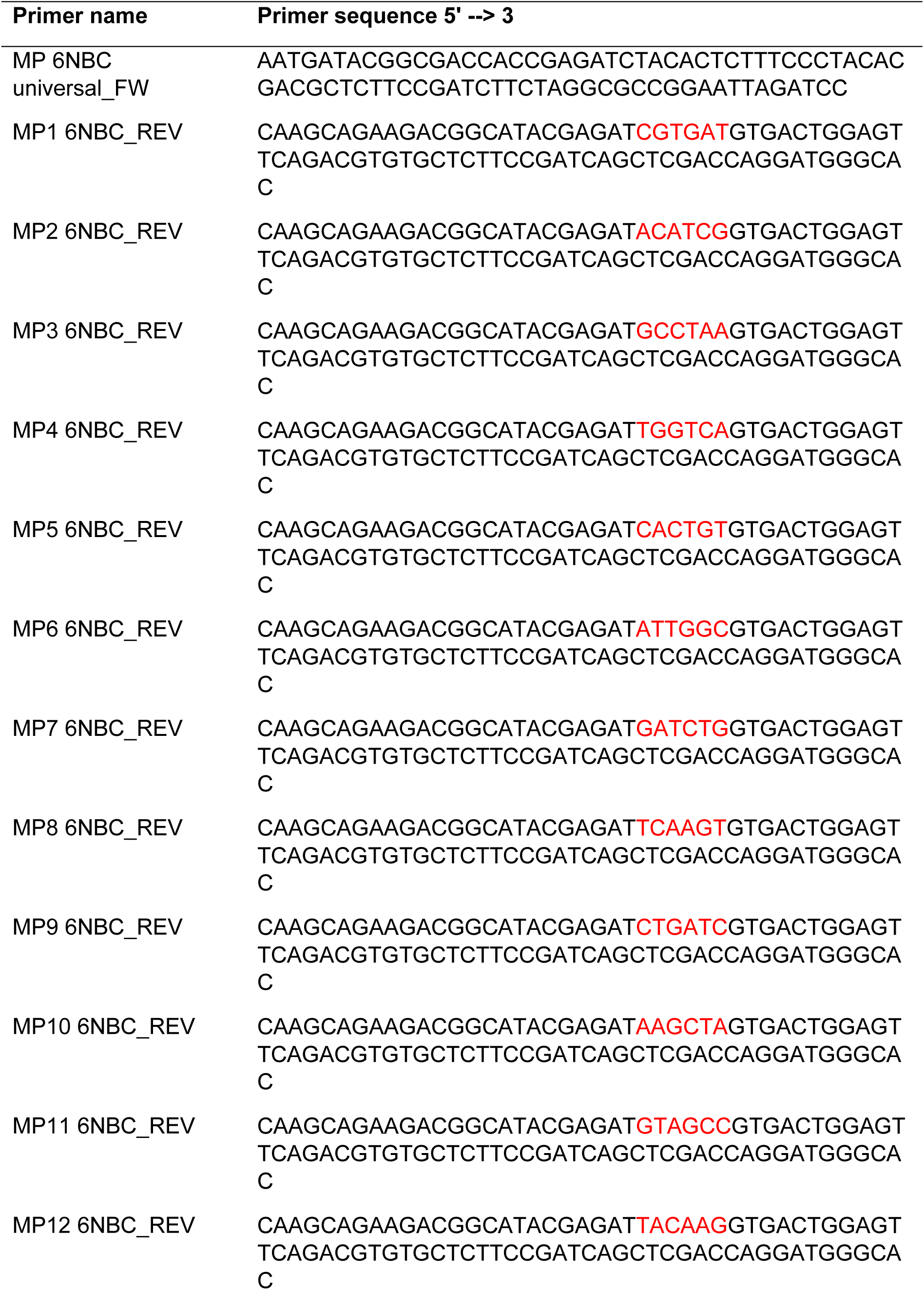
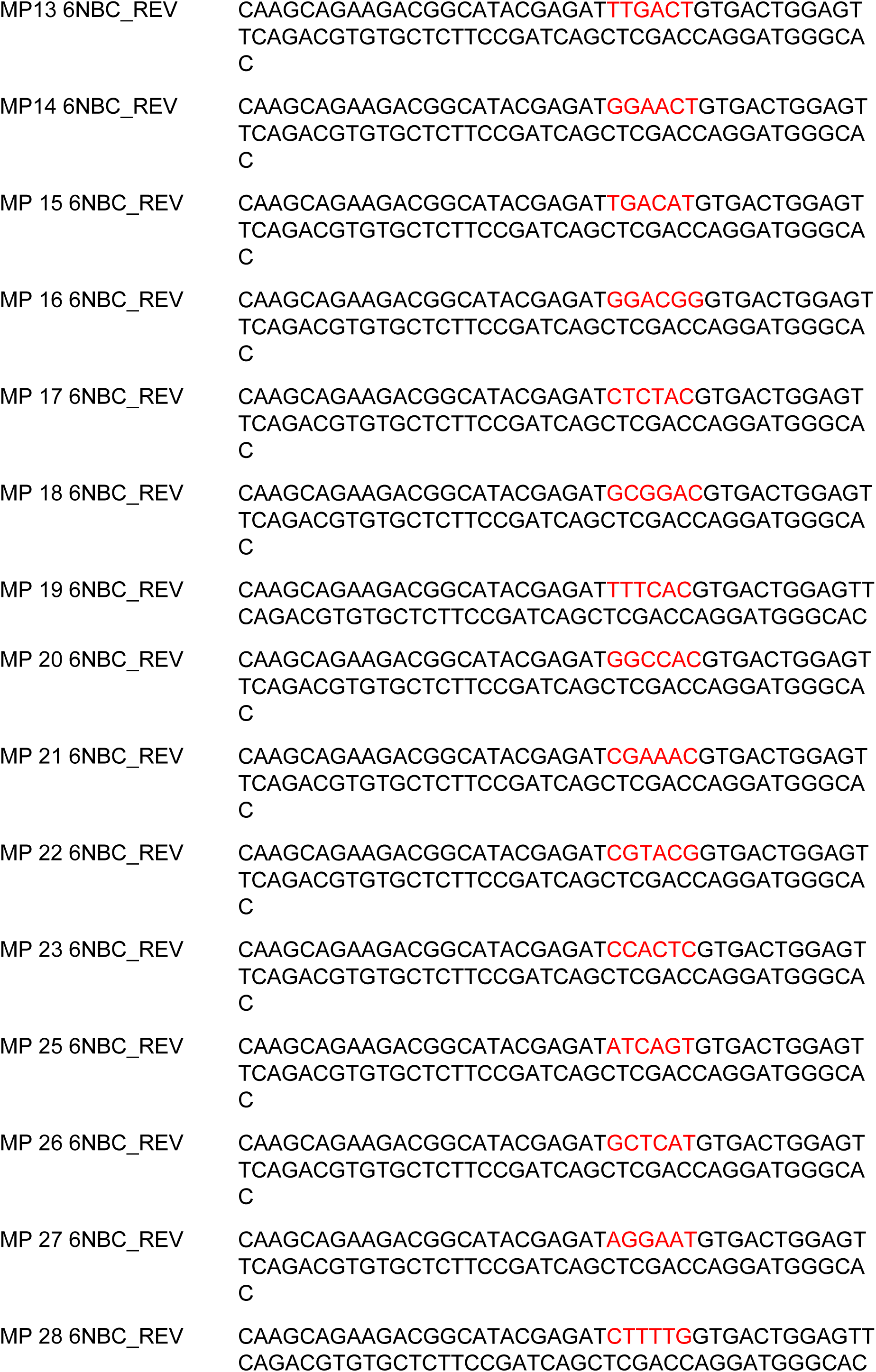
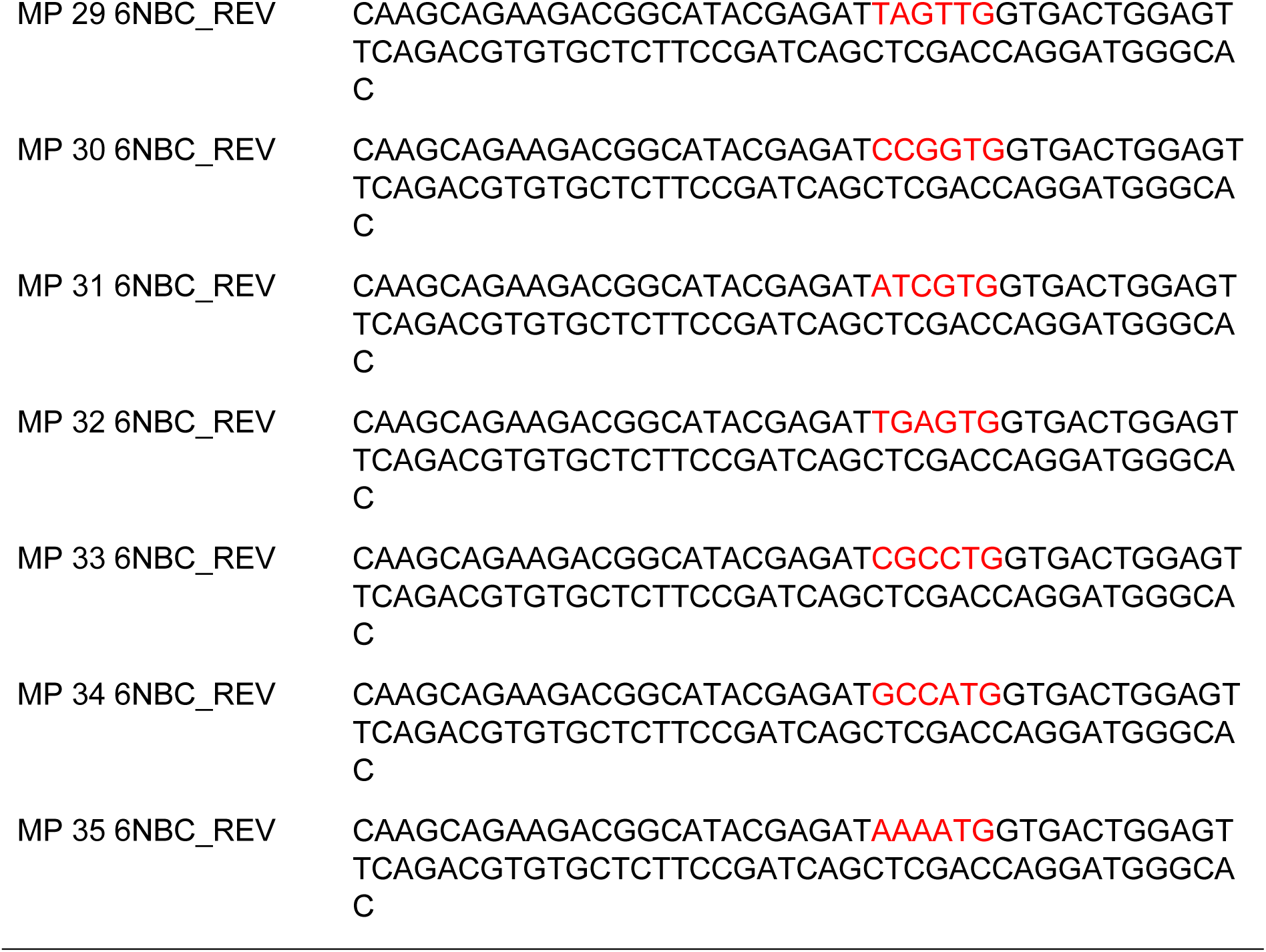

### *In vivo* experiments

All animal experiments were performed in accordance with and approved by Landesamt für Verbraucherschutz und Ernährung (LAVE), formerly LANUV (Landesamt für Natur, Umwelt und Verbraucherschutz Nordrhein-Westfalen). For all *in vivo* experiments, both male and female NMRI nude mice were divided between all experimental groups. All animal experiments and data analyses were performed without blinding and no power analysis was conducted to pre-determine group size. Mice were housed under specific pathogen-free conditions with controlled temperature, humidity and a 12-hour light/dark cycle. Food and water were provided ad libitum.

### Subcutaneous tumor injection

Tumor cells were subcutaneously injected into the flanks of recipient mice to study tumor formation, growth and therapeutic interventions. For this, tumor cells were cultured and harvested as described. Cells were resuspended in PBS, counted using the Countess II automated cell counter (Thermo Fisher Scientific), cell count was adjusted, and the suspension was mixed with Matrigel (Corning) at a ratio of 1:1. A 27 gauge needle attached to a sterile syringe was used for subcutaneous injection of 1×10^7^ cells at a final volume of 200 μL per injection site. Post-injection tumor growth was monitored using calipers and tumor volume was calculated using the formula V = (length × width^2^) × 0.5. Treatments were started upon establishment of palpable tumors as indicated below. At this point, one mouse was sacrifized to assess transplantation/growth effects of each cell line (t_0_). At the end of the experiment, mice were sacrificed using approved protocols to minimize pain and distress, and tumors were collected for further analysis.

### *In vivo* drug treatment

All drugs were applied by oral gavage once daily (qd) in the following concentrations: Imatinib mesylate (LC-Labs I-5508, CID: 123596) 200 mg/kg in 5 % glucose, sunitinib malate (LC-Labs S-8803, CID: 6456015) 40 mg/kg in NaCl/ 0.4 % Tween80 + 0.5 % carboxymethylcellulose, and trametinib (LC-Labs T-8123, CID: 11707110) 3 mg/kg in corn oil + 4 % DMSO.

### *In vivo* tumor digestion for DNA isolation and barcode sequencing

For the isolation of DNA, tumors were harvested and subsequently enzymatically digested. For this, the tumor was minced into small pieces and then incubated in digestion medium (70 % HBSS (w/o Ca^2+^, Mg^2+^), 10 % collagenase IV, 10 % dispase, 10 % 0.25 % trypsin) at 37 °C in a hybridization oven rotating at full speed for 30 min. Afterwards, an ice-cold quench solution (90 % Leibovitz’s L-15, 10 % FBS, 0.375 % DNase) was added. The suspension was filtered with a 40 µm cell strainer (Falcon Scientific) and centrifuged at 300 x g. DNA was then isolated with the Puregene Kits (QIAGEN) or the Monarch® Spin gDNA Extraction Kit (New England Biolabs) according to the manufacturer’s instructions. Then, barcode amplification and sequencing were performed as described above.

### Statistics

Tumor volume and cell growth and confluency graphs were generated using the GraphPad Prism software (GraphPad Prism, RRID:SCR_002798).

### Modeling of quantitative treatment resistance (QTR)

To quantify genotype-specific drug responses in mixed, DNA-barcoded cell populations (“pools”), we developed a hierarchical Bayesian framework that jointly models count-compositional sequencing data and independent measurements of total population size. This integration is essential because current high-throughput sequencing provides only relative clonal abundances, but the inference of clonal treatment response in a mixture of clones requires knowledge of absolute clonal expansion under treatment. By combining compositional inference with volumetric information, our framework yields a clone-specific quantitative treatment resistance (QTR) metric with an appropriate propagation of uncertainty from the two different experiments.

### Bayesian model for barcode counts

For each treatment condition *k*, barcode count vectors 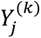 from replicate *j* were modeled using a Dirichlet–multinomial (DM) likelihood,

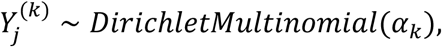

where *α_k_* = (*α_k_*_1_, …, *α_kL_*) is a vector of concentration parameters for treatment *k*. The DM distribution captures both multinomial sampling variability and biological over-dispersion across replicates.

The concentration parameters were parameterized as

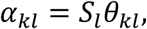

where *θ_kl_* is a compositional parameter describing the relative contribution of clone *l* under treatment *k*, and *S_l_* > 0 is a clone-specific precision parameter governing dispersion. Larger *S_l_* values concentrate observations more strongly around the expected composition, whereas smaller values allow greater variability across replicates.

The regression coefficients *β_kl_* quantify the strength of the effect of treatment *k* on clone *l*. With the following (“softmax”) transformation this effect strength is transformed into effects on the composition that we describe in terms of the previously introduced compositional parameters *θ_kl_*:

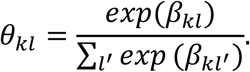

The transformation makes sure that the compositional fractions *θ_kl_* of clones *l* are all positive and sum up to one for each treatment *k*. Regression coefficients *β_kl_* were assigned weakly informative heavy-tailed priors. We used a hierarchical model with partial pooling between *β_kl_* of different clones *l*. Precision parameters *S_k_* were assigned log-normal priors to ensure positivity and the ability to model dispersion.

This model yields posterior distributions of the expected relative abundances

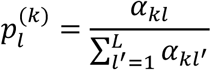

representing the inferred clonal composition under treatment *k* (after a time *l*). This final clonal composition inferred from the Bayesian model accounts for uncertainty coming from the sequencing experiment and the variability between replicates.

### Bayesian model for total population size

To translate relative abundances into absolute clonal expansion, total pool size was modeled independently for each experiment type.

For *in vivo* experiments, tumor volumes in replicates *j* under treatment *k* were modeled as

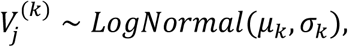

as appropriate for anatomical volumes^37^. Weakly informative priors were placed on *μ_k_* and *σ_k_* to accommodate biological heterogeneity and to regularize inference.

For *in vitro* experiments, confluency measurements were scaled to the unit interval and modeled as

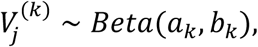

with hierarchical priors on *a_k_*, *b_k_* configured to ensure mono-modality. The volumetric measure in the previous equation is constrained to interval 0 to 1. In our experiments a confluency of 1 (or 100%) always corresponds to the same absolute area of a culture vessel, so that confluency values correspond up to a constant factor to absolute volumetric values. Note that confluency values close to saturation invalidate the assumption of approximately exponential growth underlying the equation below for the quantitative treatment resistance r_l_^(k)^ and should be avoided.

These two volumetric models yield posterior samples of the respective measure *V*^(*k*)^(*t*) at time *t*, representing the total population size under treatment *k* at that time.

### Fractional size and quantitative treatment resistance

Absolute clonal size was defined as the fractional size *ν_l_^(k)^(t)* of clone *l* under treatment *k* at time *t*,

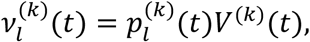

which combines compositional inference and volumetric measurement in a joint probabilistic framework.

The Quantitative Treatment Resistance (QTR) is defined as

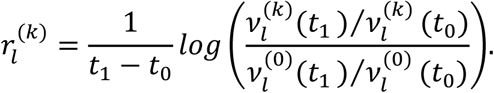

The first factor normalizes to observation time interval from *t*_0_ to *t*_1_. The logarithmic term quantifies how fractional size of clone *l* under treatment *k* changes from *t*_0_ to *t*_1_ (enumerator) in comparison to the corresponding change of fractional size under the control treatment 0. For instance, if the clone *l* grows under treatment *k* equally well to control, the argument of the logarithm is 1 and thus the quantitative treatment resistance is 0. If the argument of the logarithm is greater 1 the clone grows better under treatment than in control, QTR is positive, we have treatment resistance. Otherwise the QTR is negative, and the clone is sensitive to the treatment. As the name suggests, QTR quantifies the degree of sensitivity or resistance on a theoretical range from −∞ to ∞. QTR is given as a (posterior) probability distribution to indicate its uncertainty.

In the present work, we have simplified the equation for *r_l_^(k)^* because in the experiment we started always from the same pool in the treated and control branches so that 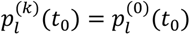, so that the corresponding factors cancel and the QTR becomes

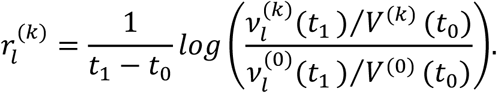

### Posterior inference and uncertainty propagation

We drew samples from the posterior with an MCMC procedure (see below) from both models, the one for the relative clone abundances and the one for the volumetric measure (tumor volume or confluency). For each posterior draw *S*, the corresponding fractional sizes were computed as

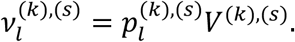

Treatment resistance samples were then obtained from:

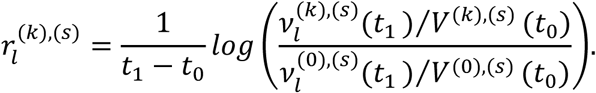

This equation fully propagates uncertainty from both sequencing and volumetric measurements into the posterior distribution of treatment resistance.

### Statistical inference

All Bayesian models were estimated in R using Stan, which performs Markov chain Monte Carlo (MCMC) sampling via the No-U-Turn Sampler (NUTS)^38, 39^. Each model was fitted using two independent chains with 10,000 iterations per chain, yielding 20,000 posterior samples. Convergence was assessed through standard diagnostics, including the absence of divergent transitions and the potential scale reduction factor (*R^*), which was required to be close to 1.0 for all parameters^40, 41^.

The tuning parameter adapt_delta, which specifies the target average acceptance probability of the NUTS proposals was set to 0.999 and the initial step size was set to 0.5. The maximum tree depth was set to 18 for the sequencing and tumor-volume models and to 12 for the confluency model.

## Data availability

The datasets generated and analyzed during the current study are available on the ENA server. The repository contains all *in vitro* and *in vivo* barcode-readout datasets for pools I, II, IIIa, and IIIb. The R package to perform all QTR analysis *in vitro* and *in vivo* is available on GitHub under the name “barmixR” https://github.com/MohammadDarbalaei/barmixR.

## Notes

https://github.com/MohammadDarbalaei/barmixR

