## Supplemental Figures 1-6 for "Quantifying Treatment Resistance in Mixtures of Gastrointestinal Stromal Tumor Cells with BARMIX"

### Extended Data Fig. 1:

**a**

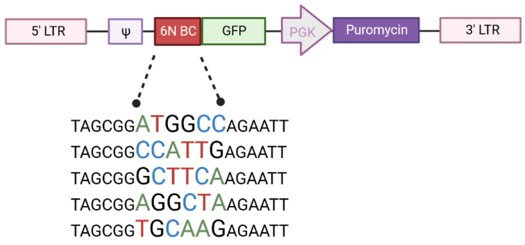


**b**

| Sub cell line | Parental T1 | TSC2 -/- | PTEN -/- | KRAS$^{G12R/G12R}$ HOM | KIT$^{A829P}$ | KIT$^{D816A}$ | KIT$^{D816E}$ | KIT$^{V654A}$ |
|---|---|---|---|---|---|---|---|---|
| BC #1 | GGCACG | GTTTAG | CTAGGT | GGGAAC | GGTCCA | CTAGAT | GGACGT / TATATA | GTAACC |
| BC #2 | TTAGAA | AGAGTT | TTTGAA | AGTTAC | TTAGAT | TGGGCC | CCAGGT | TGAGAA |
| BC #3 | CTCCAA | GAAAAA | CAGGCC | ACTCAA | ATGCAT | CAGGTC | GTGAAG | GGGCGT |

**Extended Data Fig. 1: Combination of an isogenic cell line model with molecular barcoding to analyze genotype-specific drug resistance mutations in GIST**

**a)** Schematic overview of the 6 nucleotide barcode containing vector.

**b)** Overview of the GIST isogenic cell line models and the corresponding barcode sequences. Based on the parental cell line GIST-T1, which is characterized by oncogenic activation of the KIT receptor, 7 additional cell lines with secondary resistance mutations were used to reflect the heterogeneity of therapy resistance. The parental GIST-T1 and each sub-cell line were labeled individually in three independent replicates with one distinct barcode each as indicated to generate 24 barcoded cell lines in total, representing each genotype in triplicate.

### Extended Data Fig. 2:

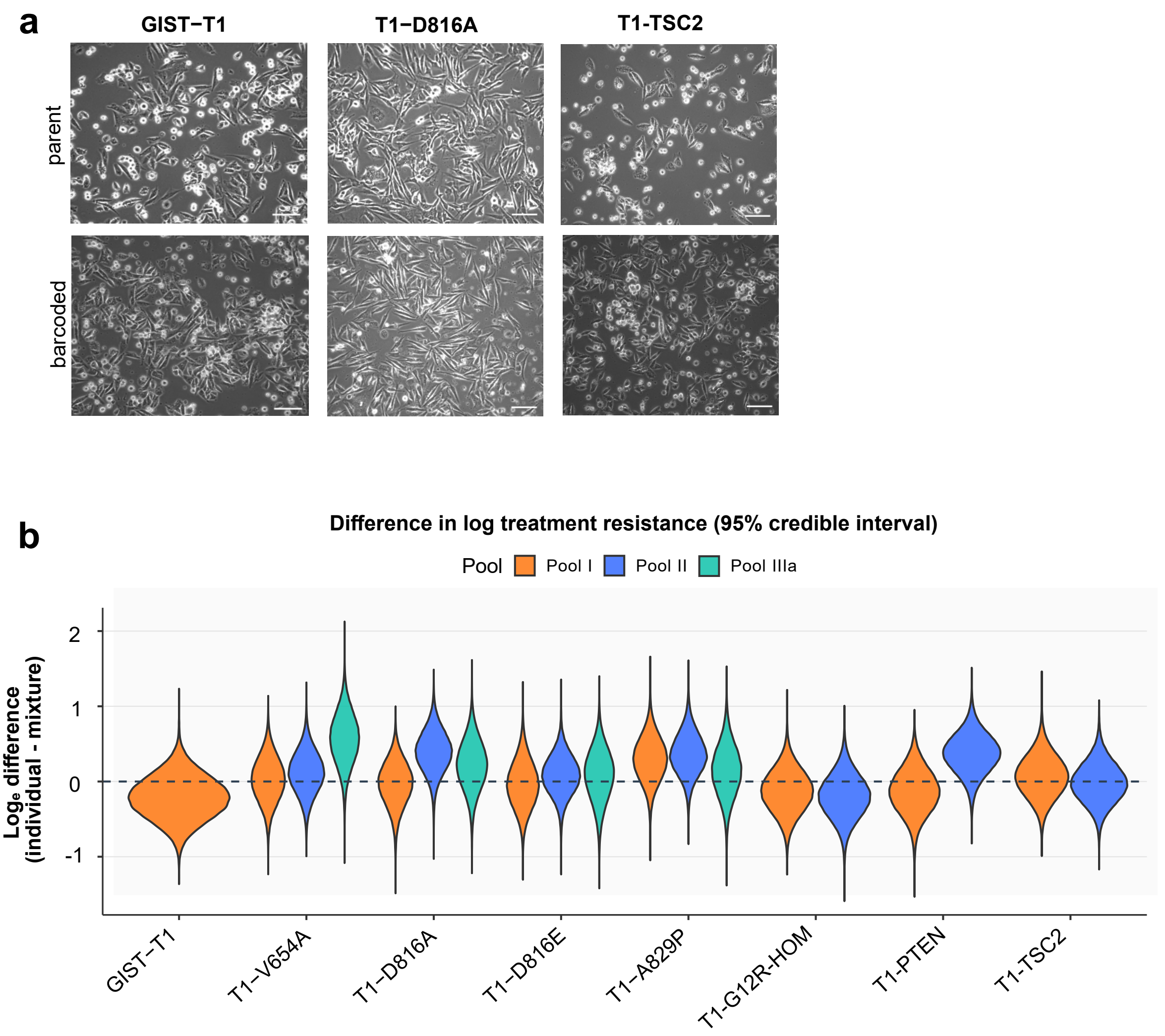

**Extended Data Fig. 2: Comparison of barcoded and non-barcoded cells with and without treatment**

**a)** Brightfield microscopic images of example cell lines. Unlabeled cell line (upper panel) and the corresponding barcoded cell line (lower panel) have similar appearance. Scale bars equal 100 µm.

**b)** Posterior distributions of $\Delta_l$=$r_{l,ind}$-$r_{l,pool}$(differences in treatment resistance between individual and pooled assays) for each cell line. Colors indicate pools. The dashed line at zero corresponds to no difference between individual and pooled assays; values above (below) zero indicate higher resistance in the individual (pooled) assay.

#### Extended Data Fig. 3 - page 1/6

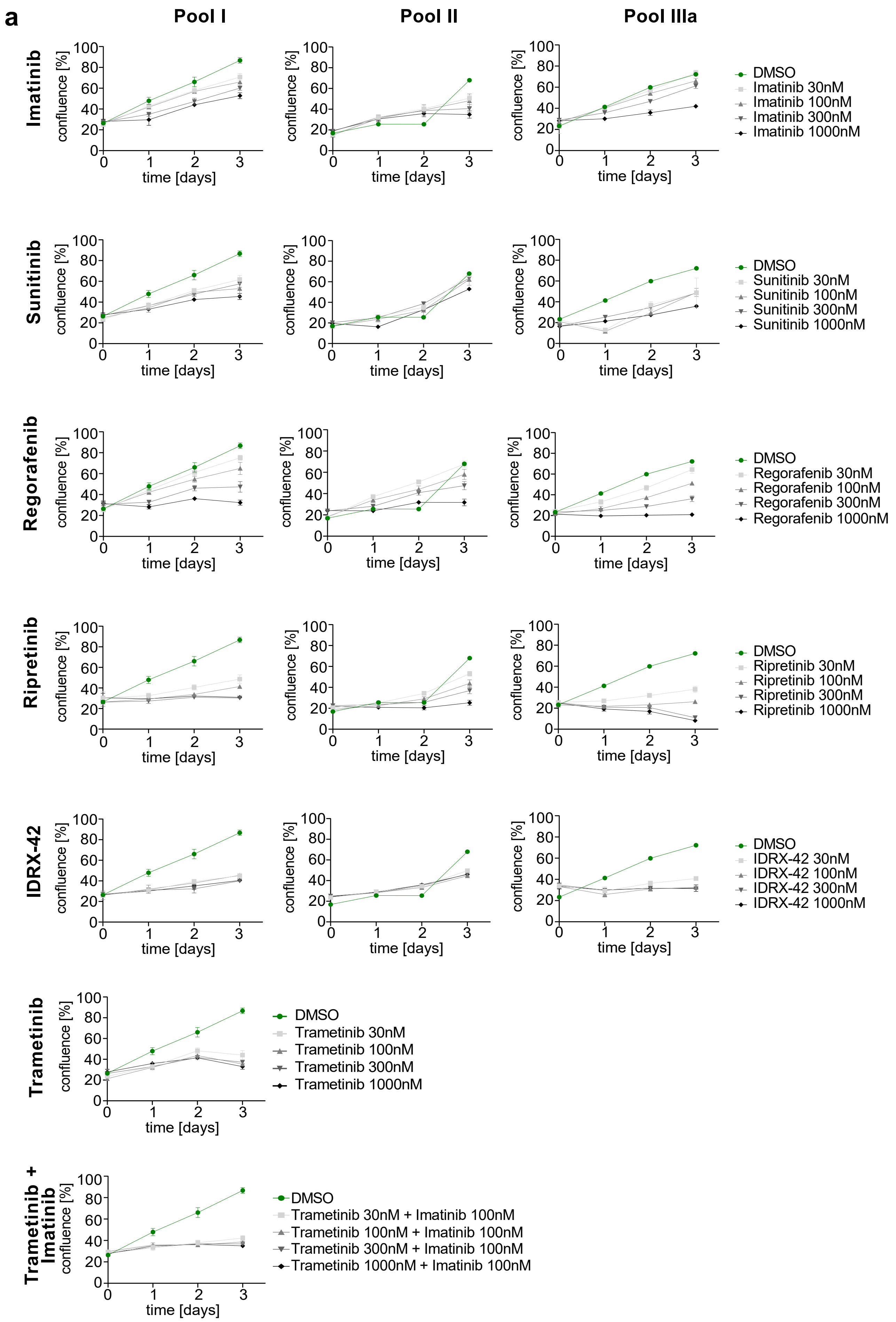

### Extended Data Fig. 3 - page 2/6

**b**

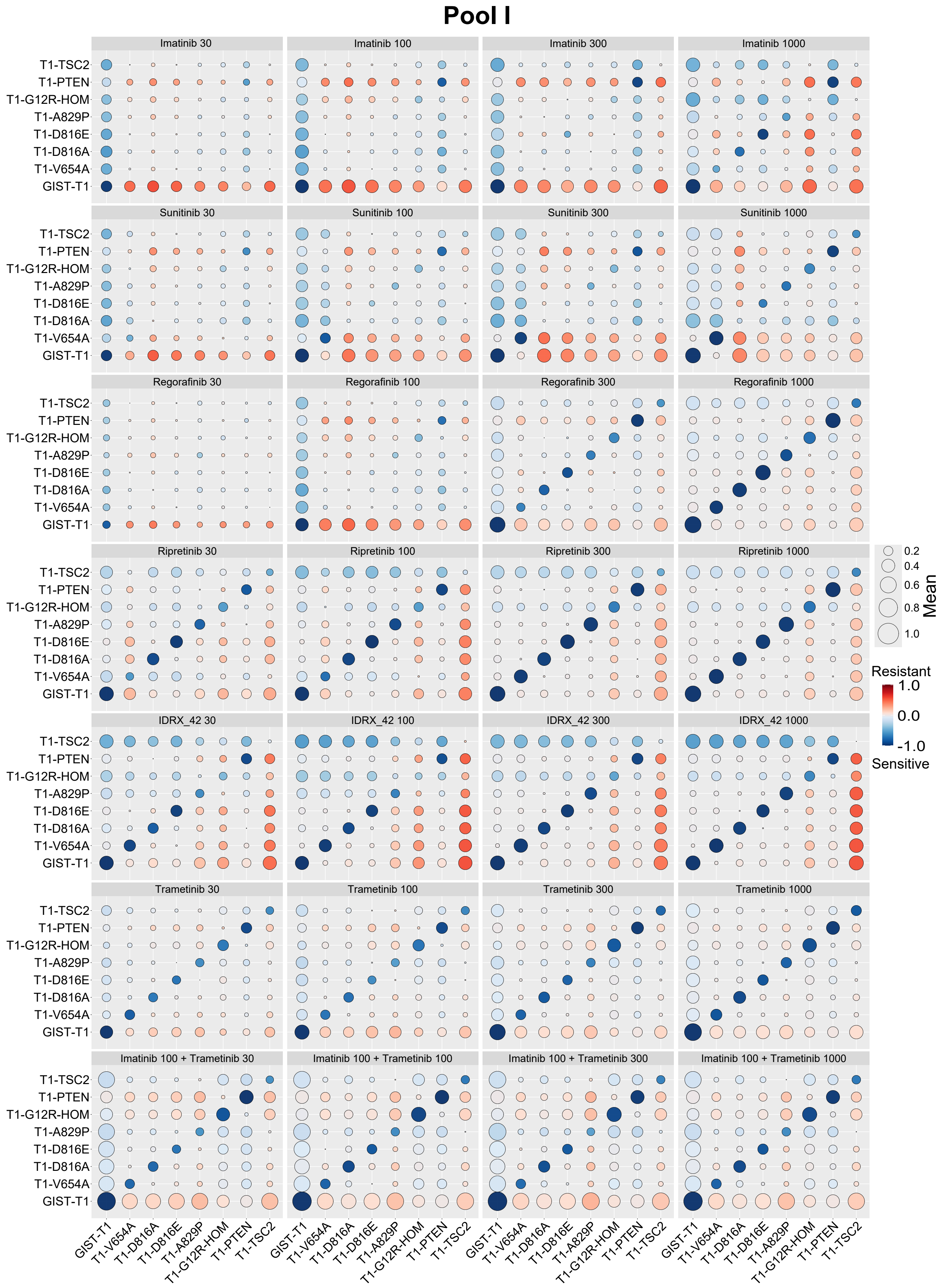

### Extended Data Fig. 3 - page 3/6

##### *in vitro* 3 day data for pool II

c

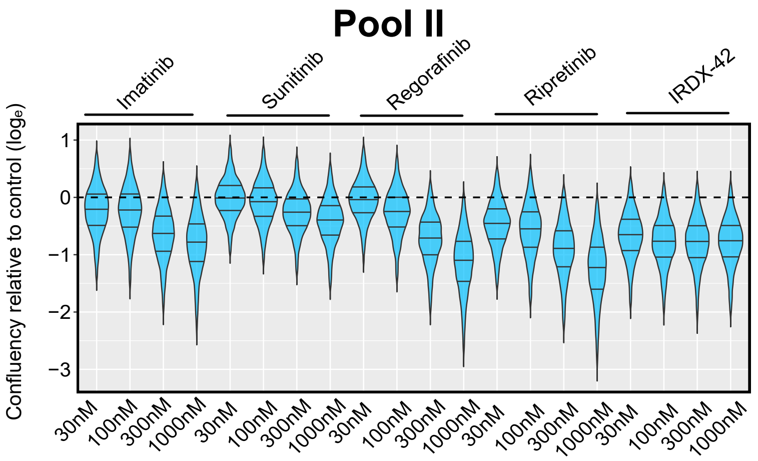


d

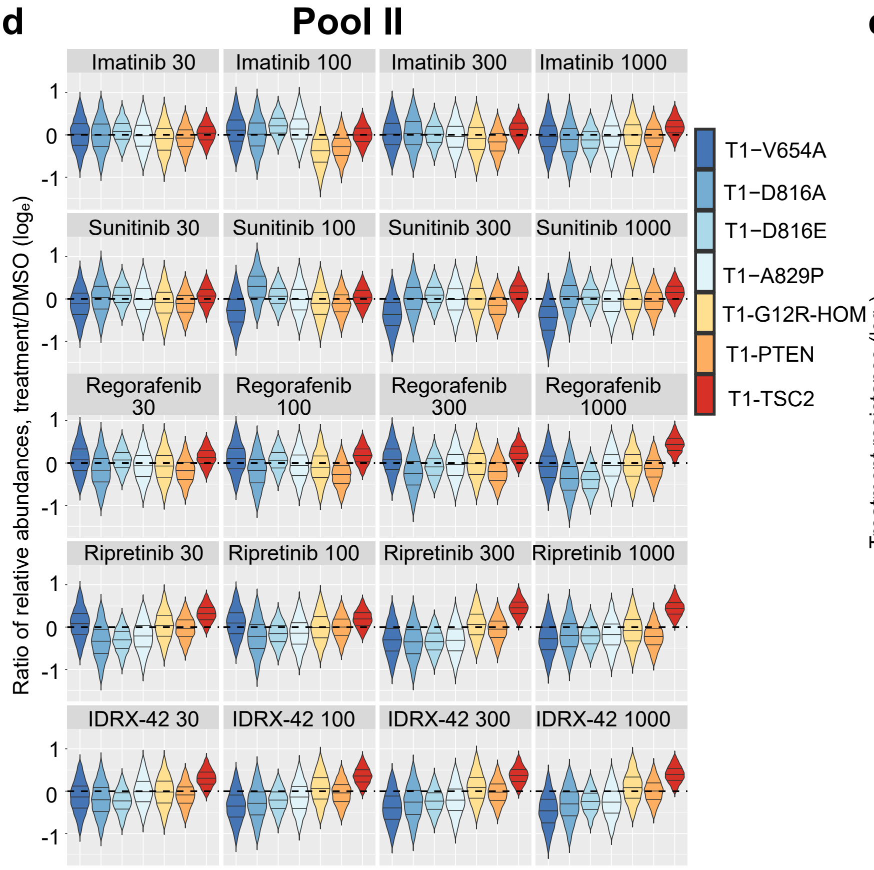


e

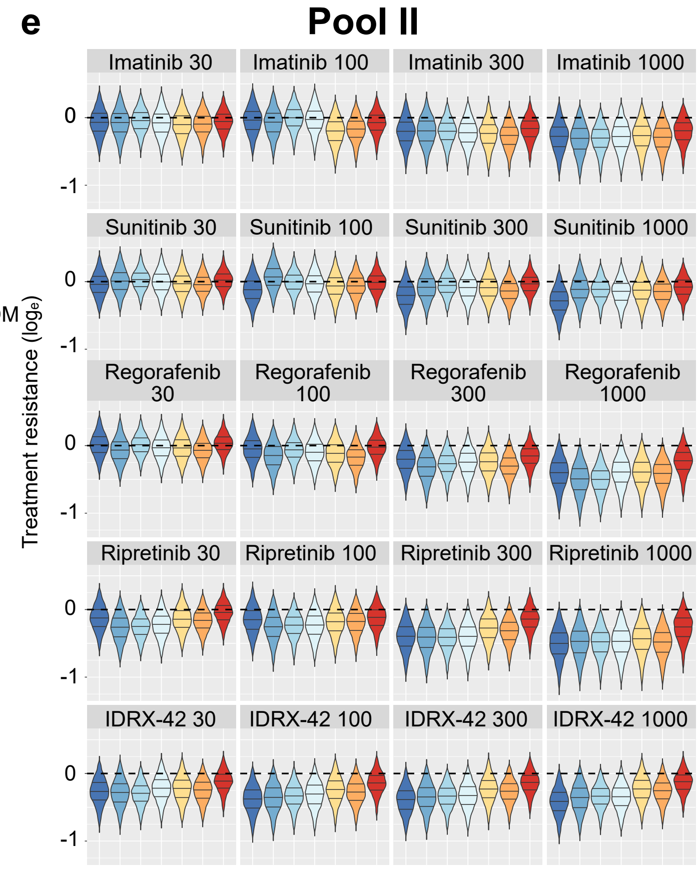


f

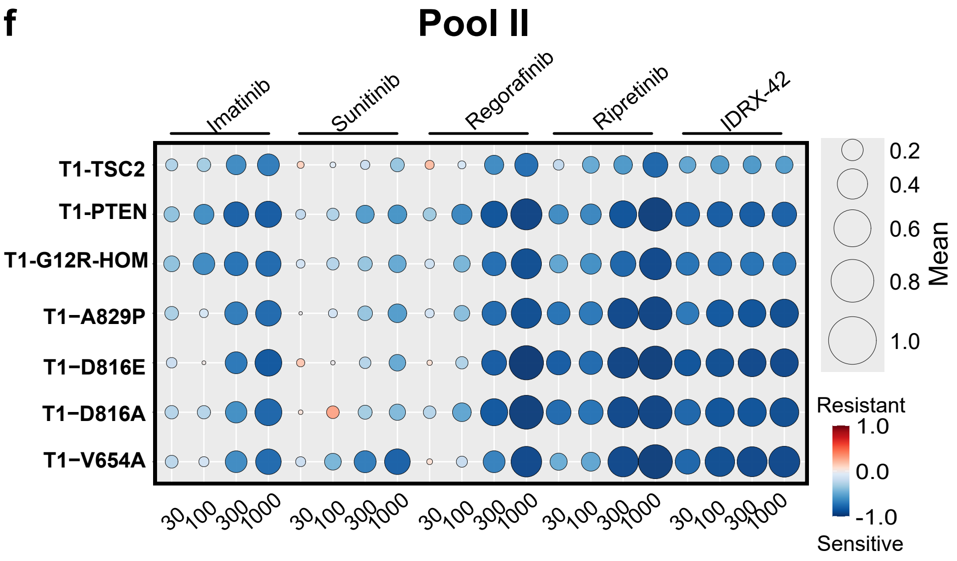

### Extended Data Fig. 3 - page 4/6

***in vitro* 3 day data for pool II**

**g**

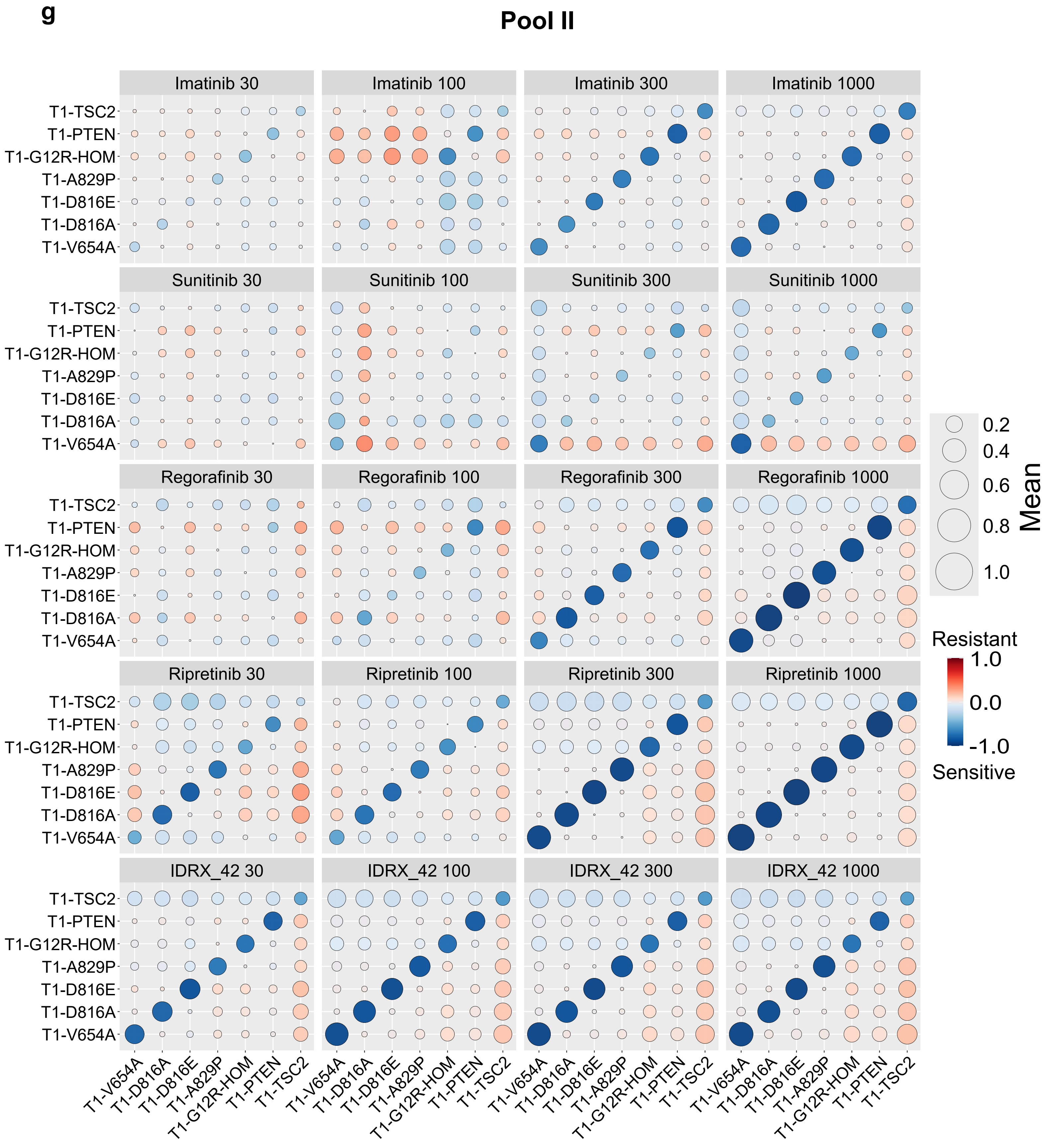

### Extended Data Fig. 3 - page 5/6

***in vitro* 3 day data for pool III**

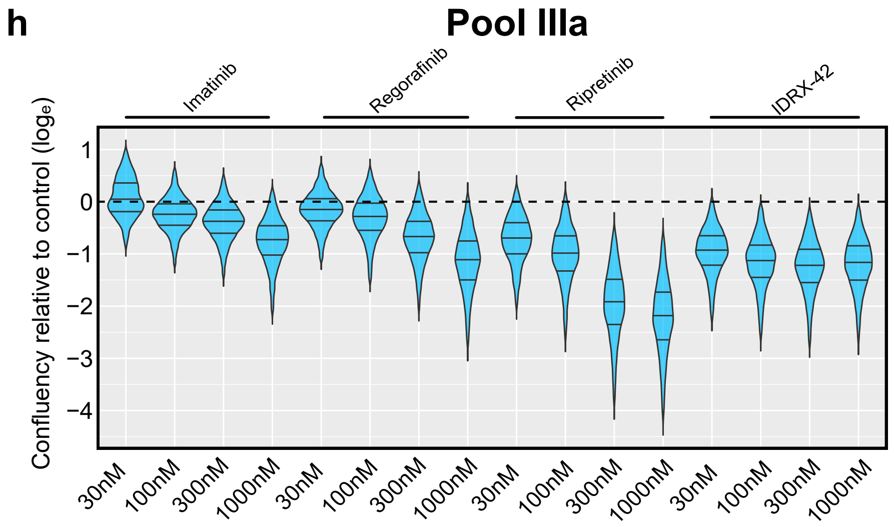


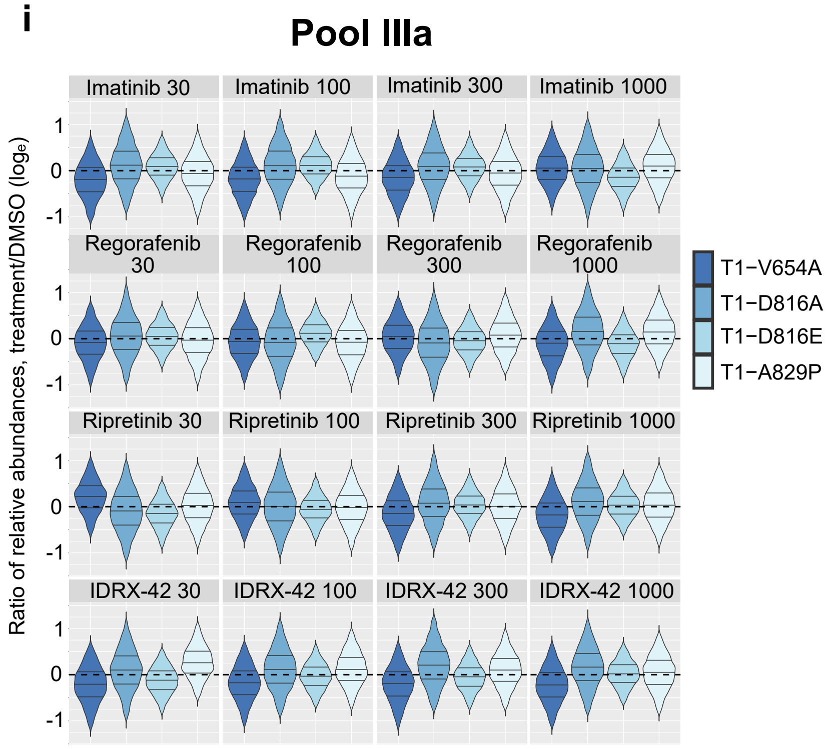


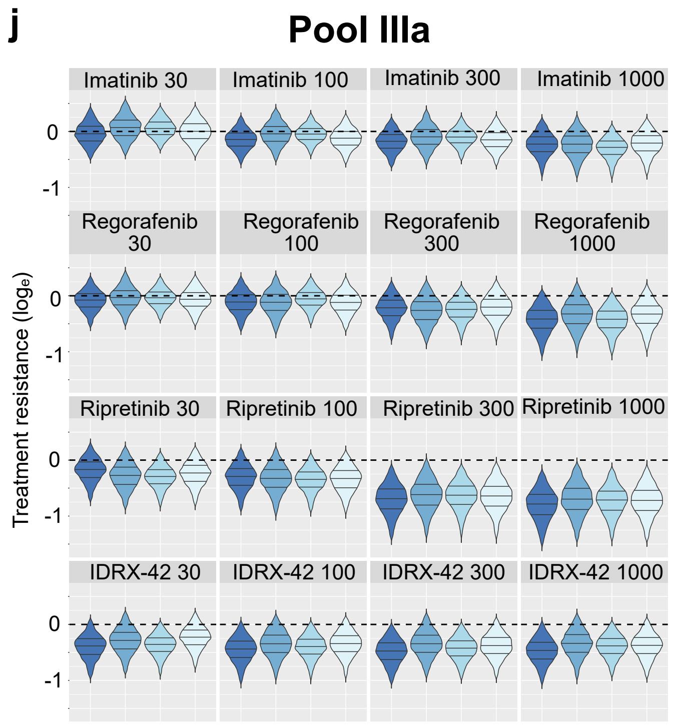


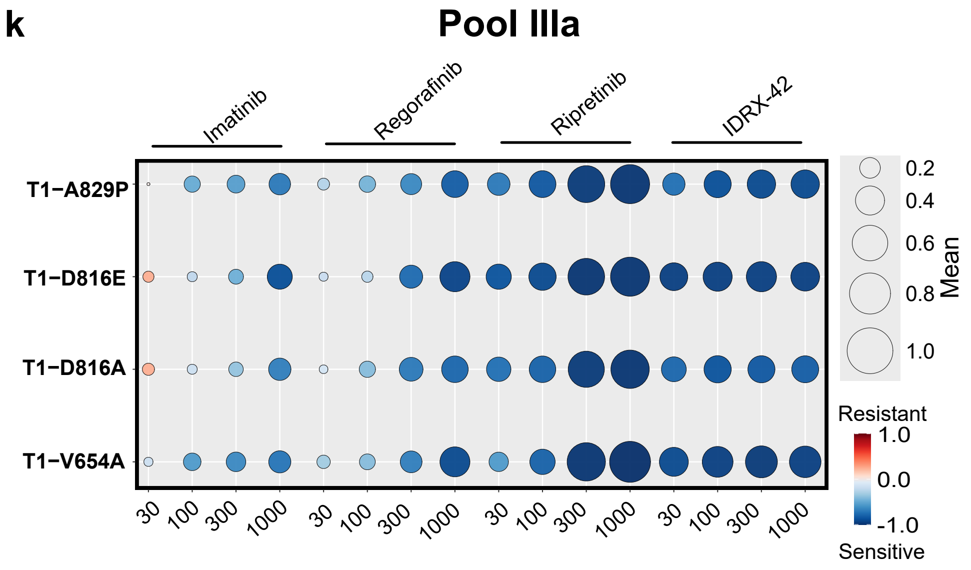

### Extended Data Fig. 3 - page 6/6

***in vitro* 3 day data for pool III**

**I**

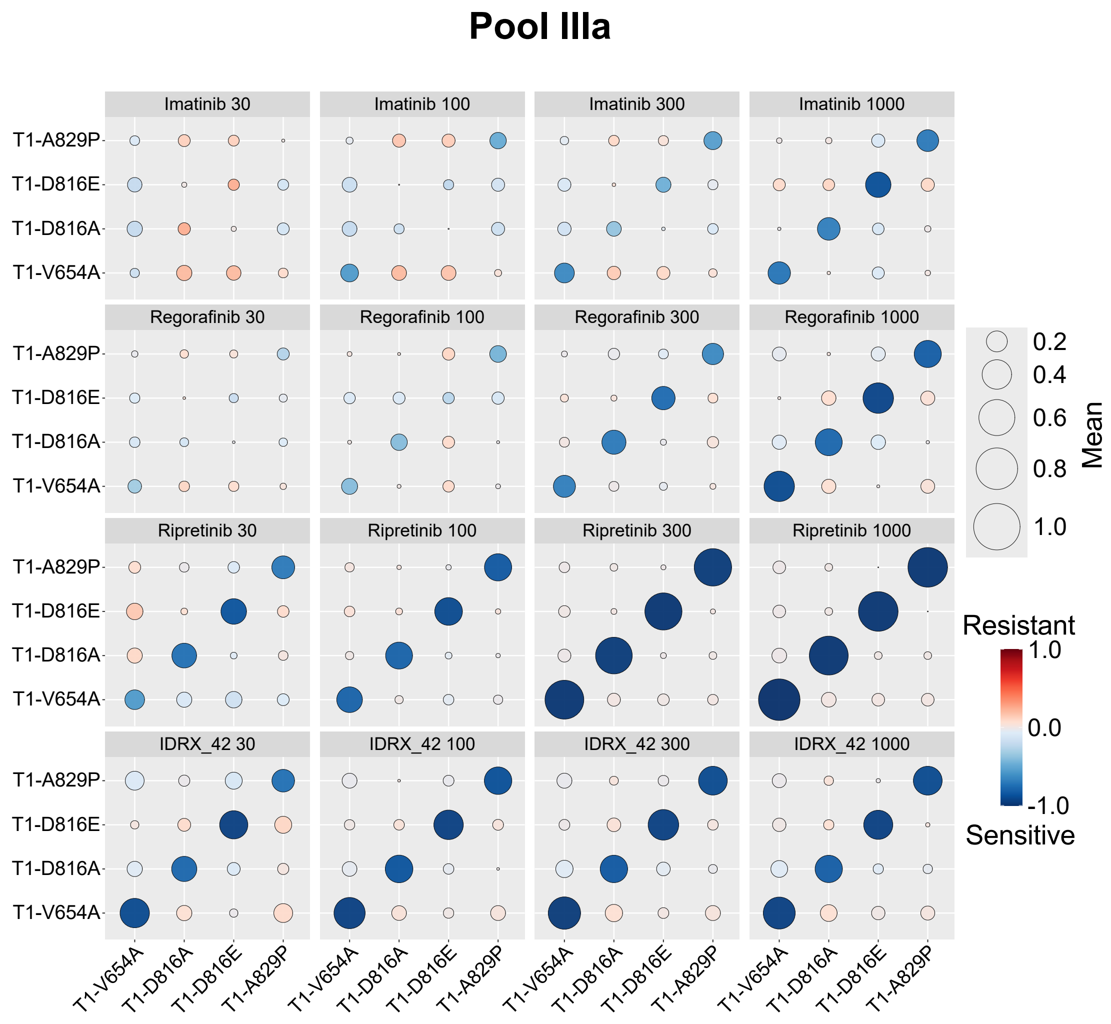

**Extended Data Fig. 3: Quantitative assessment of *in vitro* short-term treatment responses**

**a**) Relative growth of barcoded cell line pools under control (DMSO, green symbols) or experimental (black symbols) treatment over time measured by relative confluency in triplicate wells. In case error bars are not visible, they are smaller than the symbols. Treatment and concentrations as indicated.

**b)** Bubble plots depict pairwise comparisons of quantitative treatment resistance (QTR) among cell lines across individual treatment conditions in Pool I after 72 hours. Each panel represents one treatment at the concentration (in nM) indicated in the panel head, with cell lines shown on both axes. Bubble size reflects the magnitude of the *difference* of posteriors means of QTR between two cell lines, whereas diagonal bubbles are the posterior mean of QTR for each cell line. Bubble colour encodes the direction of the QTR difference (color legend on the right): red indicates that the row cell line is less resistant (more sensitive) than the column cell line, e.g. in the top left panel the bottom row of red dots means that the parental GIST-T1 cell line is less resistant against Imatinib (30nM) than all other cell lines. Consequently, the bold blue dots in the first column indicate higher resistance to 30nM Imatinib of each non-GIST-T1 cell line than GIST-T1.

**c-g) *In vitro* analysis of treatment responses in Pool II at 72 hours – compare Main Fig.3. c)** Natural-log scaled ratio of confluency between treated and control conditions, with uncertainty inferred from a Bayesian model of confluency measurements. The horizontal black dashed line marks the baseline of no change of confluency under treatment relative to control. Treatments along top, concentrations along bottom. **d)** Natural-log scaled ratio treatment to control of relative abundances, with uncertainty inferred from a Bayesian model of barcode abundance. Drug concentrations in panel heads in nM. **e)** Treatment resistance inferred by integrating Bayesian models of barcode abundance and cell confluency, with uncertainty propagated from both components. Each distribution reflects resistance on a natural-log scale for a given genotype–treatment pair. The horizontal black dashed line separates sensitive (below) and resistant (above) regimes. Drug concentrations (panel heads) in nM. **f)** Summary of treatment resistance across genotypes and drug conditions. Bubble size reflects the mean QTR; color represents direction: blue = sensitive, red = resistant. Drug concentrations in nM. **g)** Bubble heatmaps (as in Extended Data Fig. 3a) are pairwise comparisons of treatment resistance among cell lines across individual treatment conditions in Pool II after 72 hours.

**h-l) *In vitro* analysis of treatment responses in Pool IIIa at 72 hours.** Parts h)-k) as in main Fig. 3b)-e). Part l) as in Extended Data Fig. 3b).

### Extended Data Fig. 4 - page 1/3

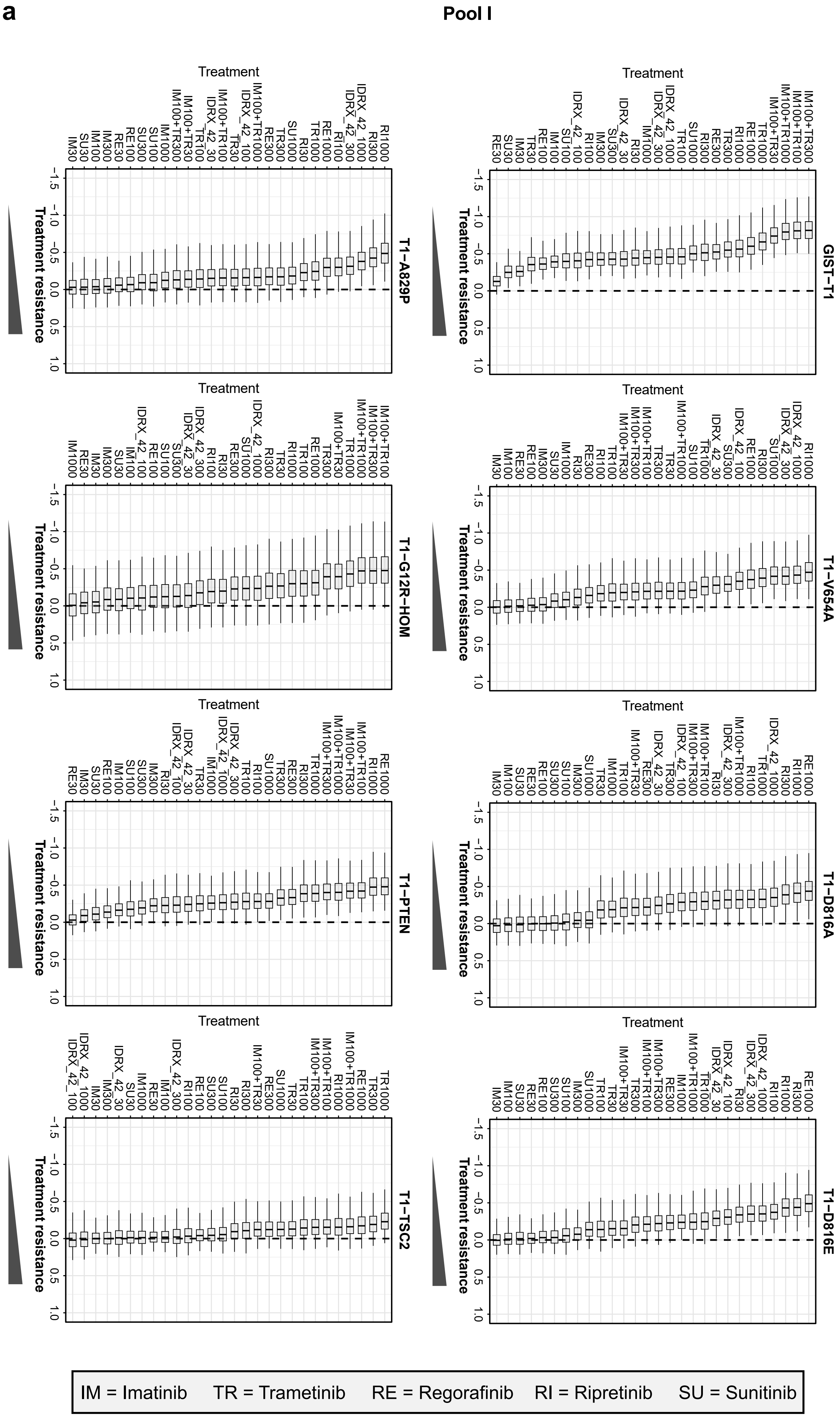

### Extended Data Fig. 4 - page 2/3

**b** **Pool II**

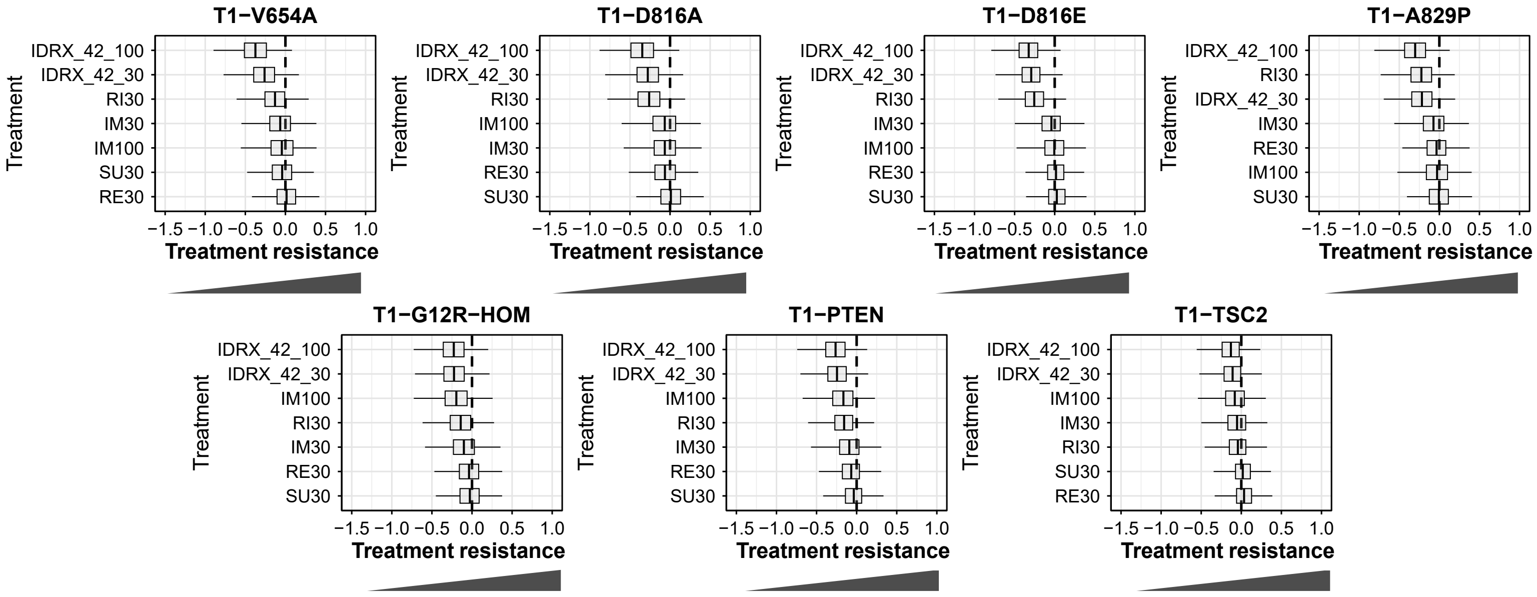


IM = Imatinib  RE = Regorafinib  RI = Ripretinib  SU = Sunitinib

**c** **Pool II**

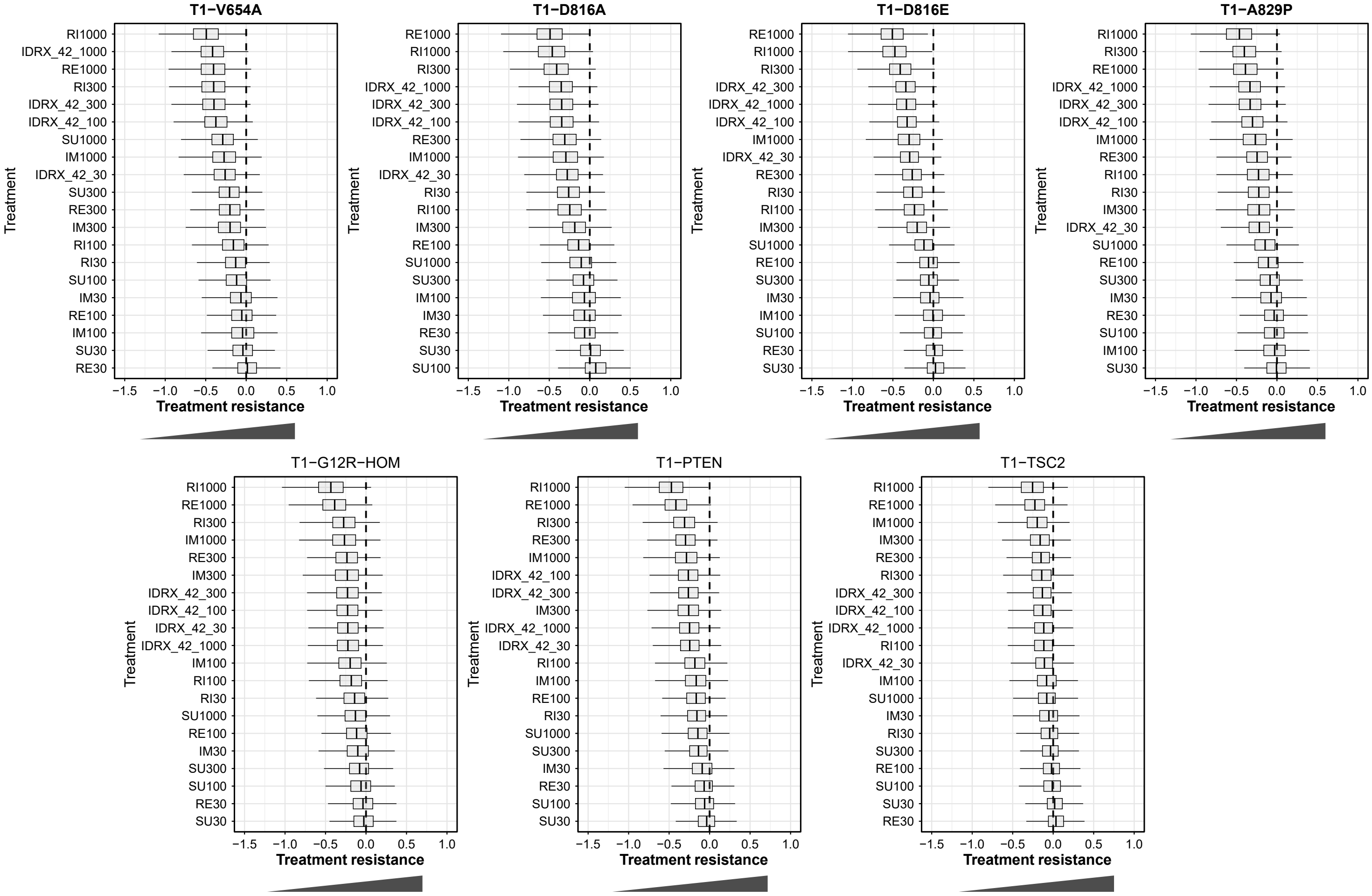

### Extended Data Fig. 4 - page 3/3

**d** **Pool IIIa**

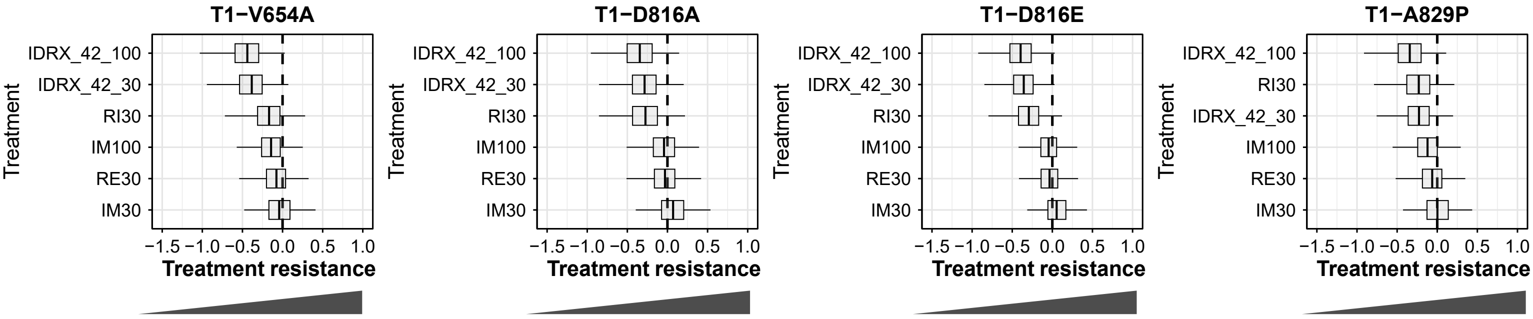


IM = Imatinib RE = Regorafinib RI = Ripretinib

**e** **Pool IIIa**

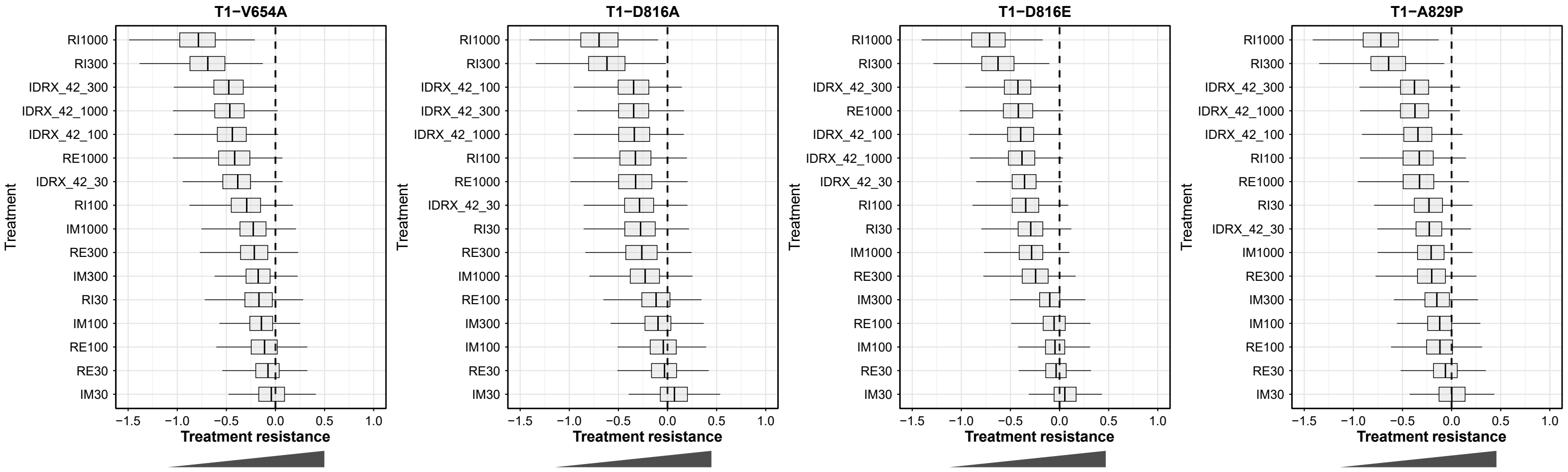

**Extended Data Fig. 4: Quantitative ranking system to support preclinical decision making**

**a)** All treatments and concentrations evaluated in Pool I after 72h hour treatment *in vitro* ranked according mean QTR as in Main Fig. 4.

**b-c)** Ranked QTRs in Pool II after 72h hour treatment *in vitro* – compare Main Fig. 4. **b)** Selected treatment concentrations reflect clinically relevant or commonly used doses. **c)** All evaluated treatments with their drug concentrations are shown.

**d-e)** Ranked QTRs in Pool IIIa after 72h hour treatment *in vitro* – compare Main Fig. 4. **d)** Selected treatment concentrations reflect clinically relevant or commonly used doses. **e)** All evaluated treatments and drug concentrations are shown.

#### Extended Data Fig. 5 - page 1/7

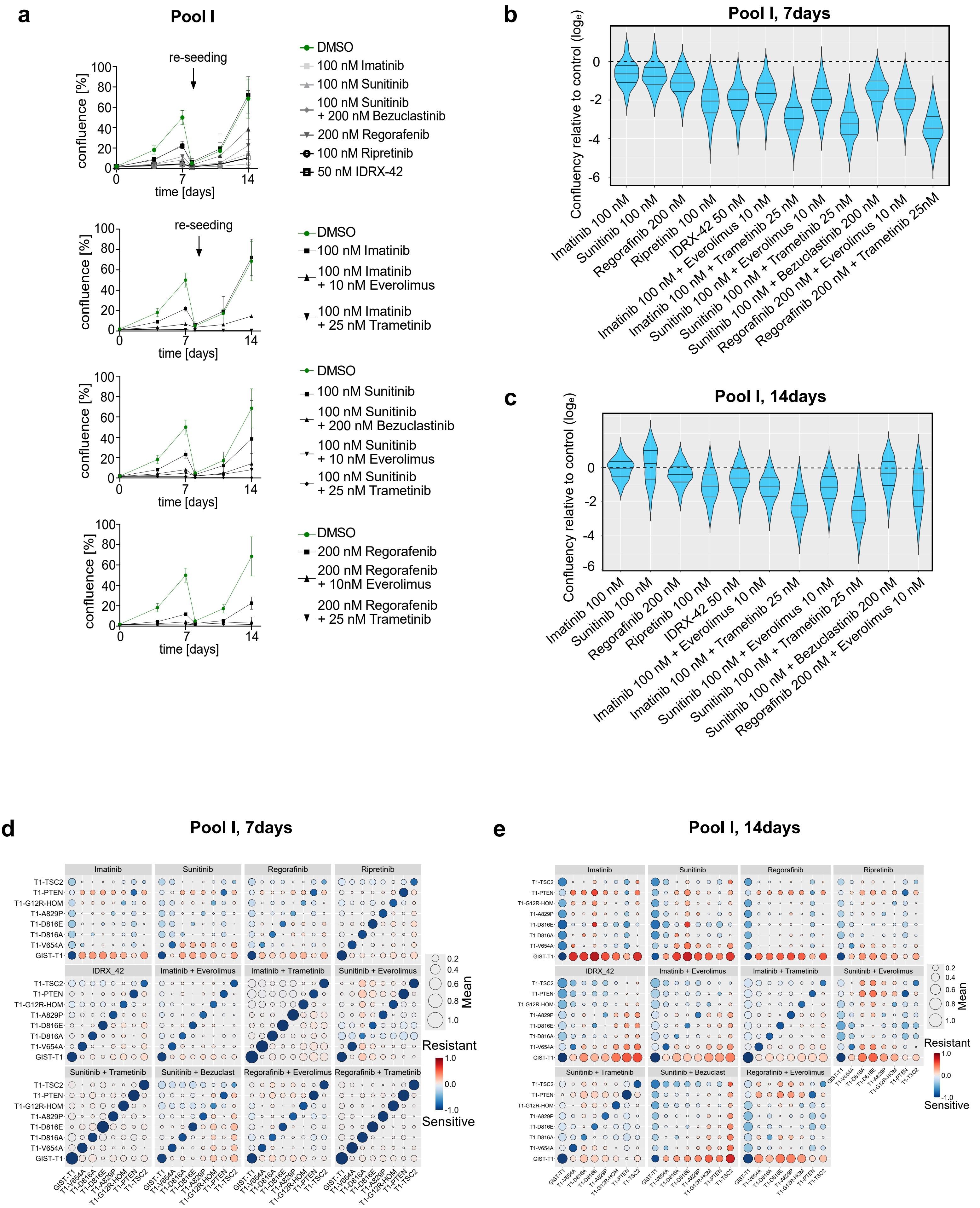

#### Extended Data Fig. 5 - page 2/7

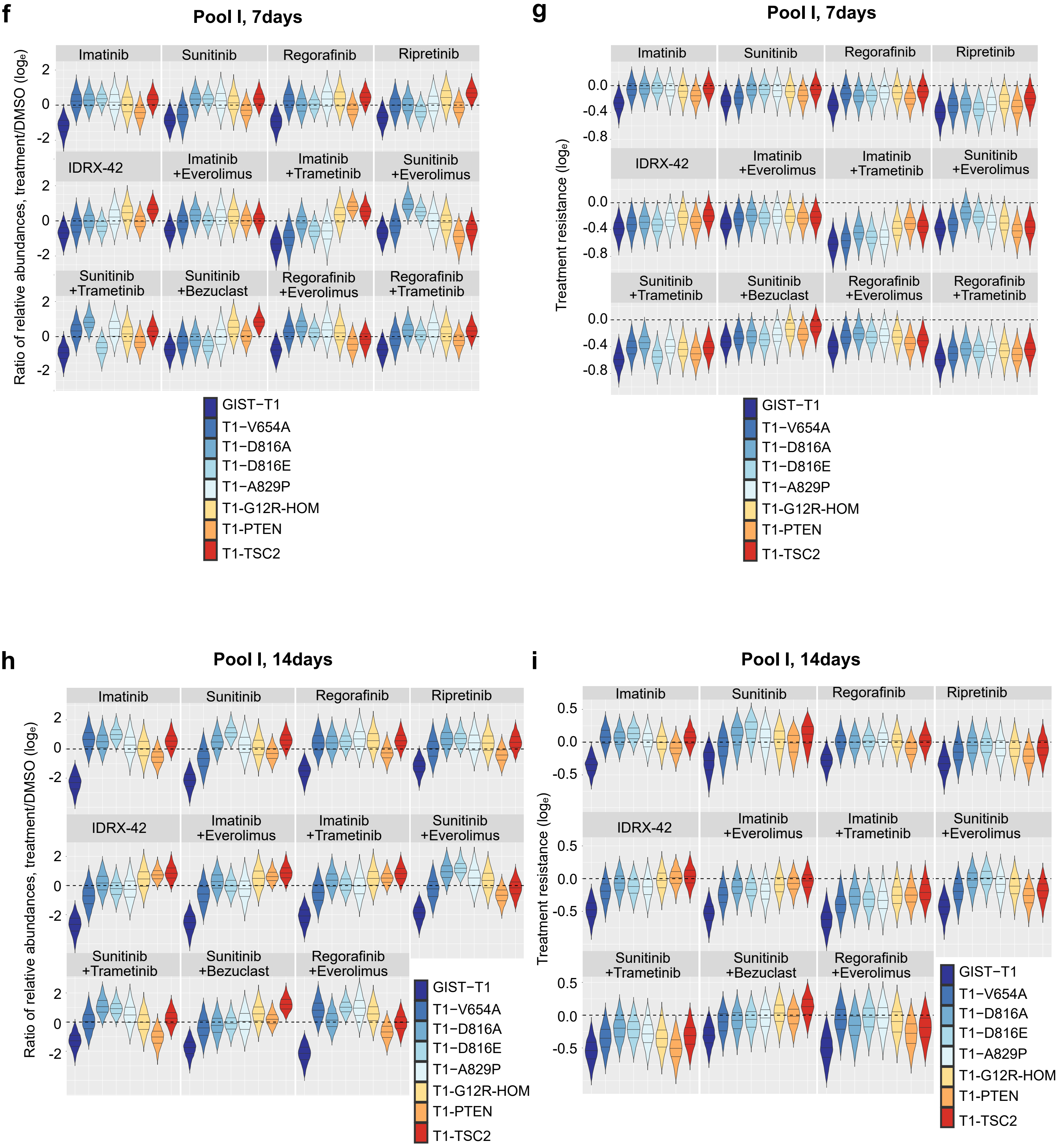

### Extended Data Fig. 5 - page 3/7

#### Pool I, 7days

**j**

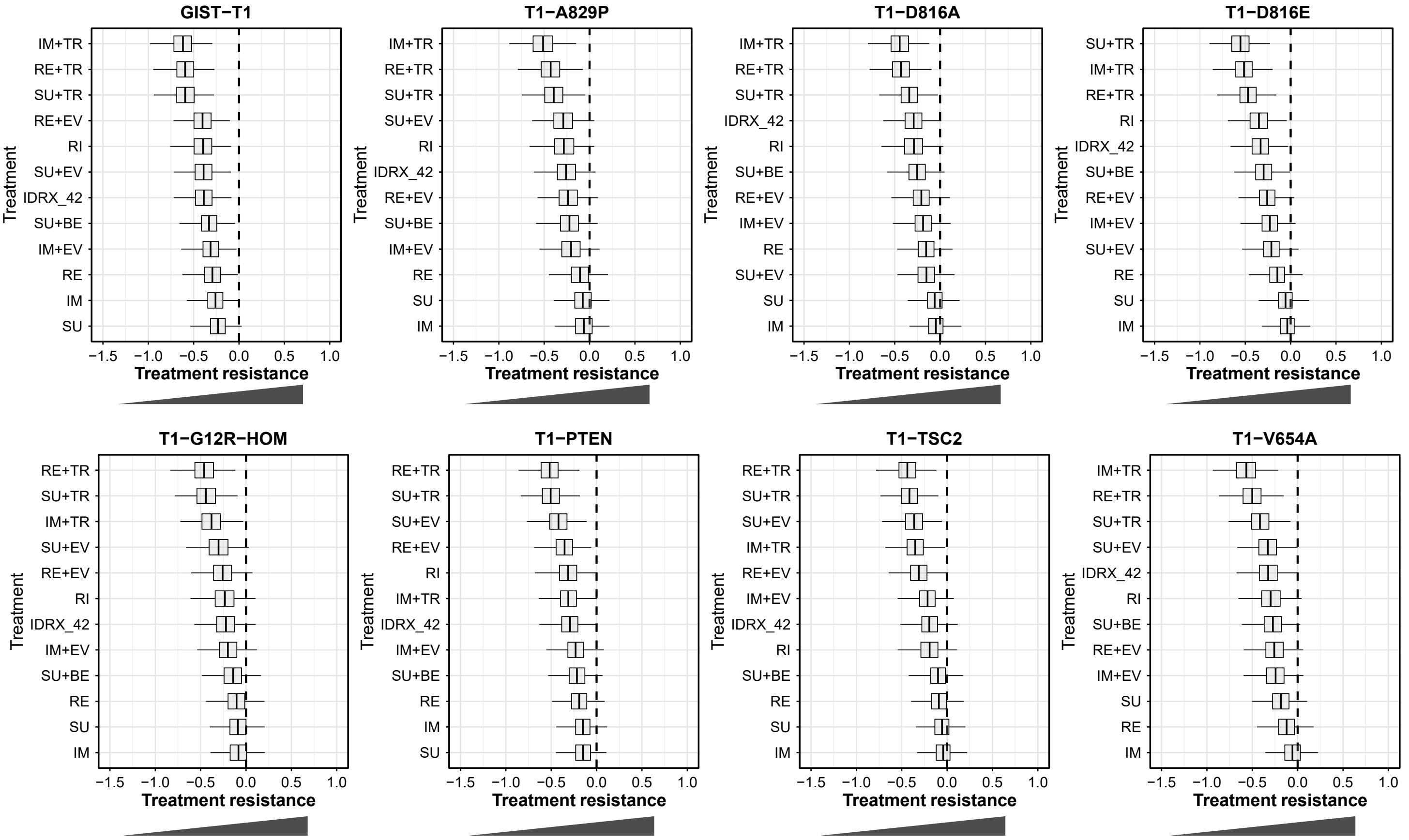


IM = Imatinib TR = Trametinib RE = Regorafinib RI = Ripretinib SU = Sunitinib EV = Everolimus BE = Bezuclast

### Extended Data Fig. 5 - page 4/7

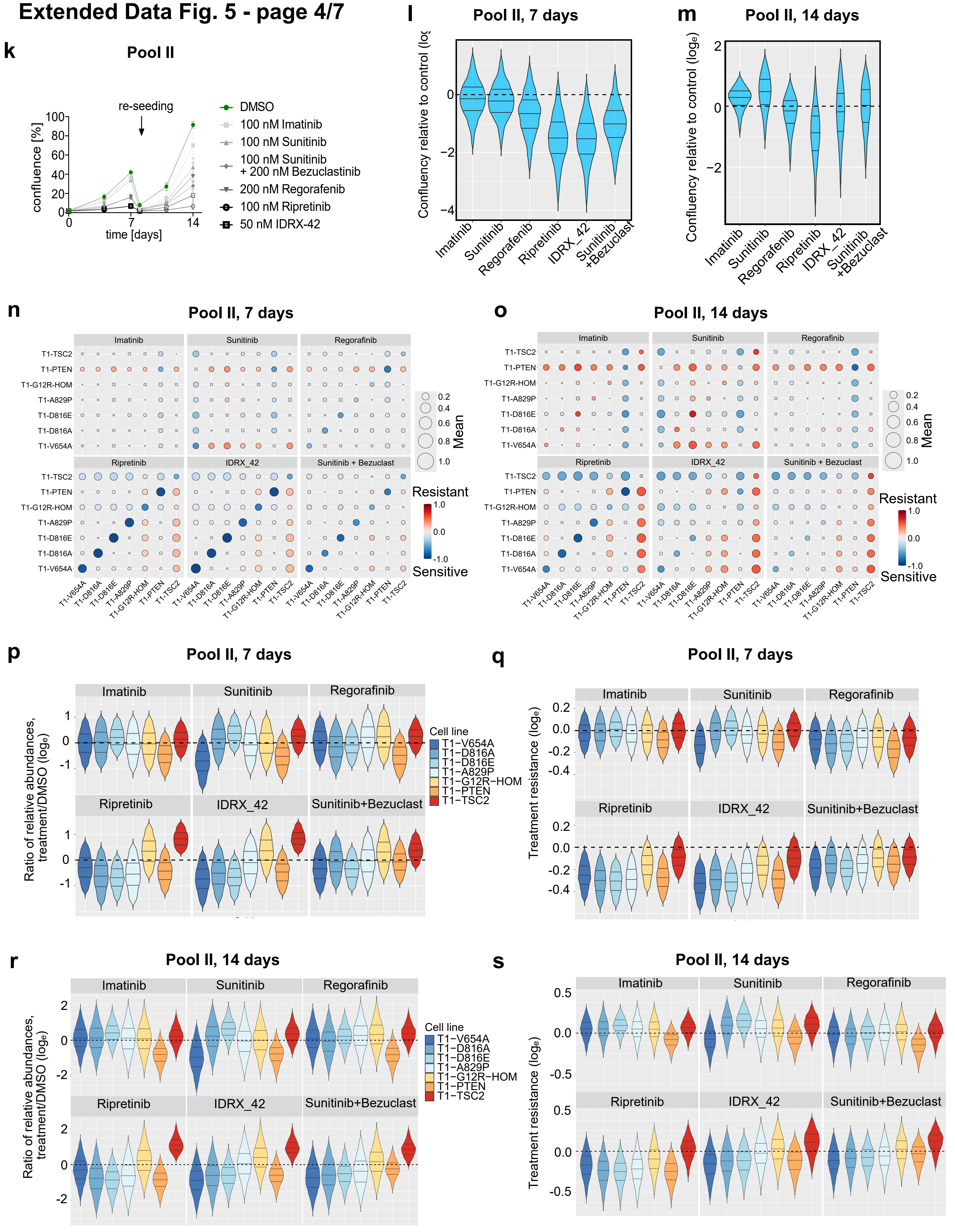

### Extended Data Fig. 5 - page 5/7

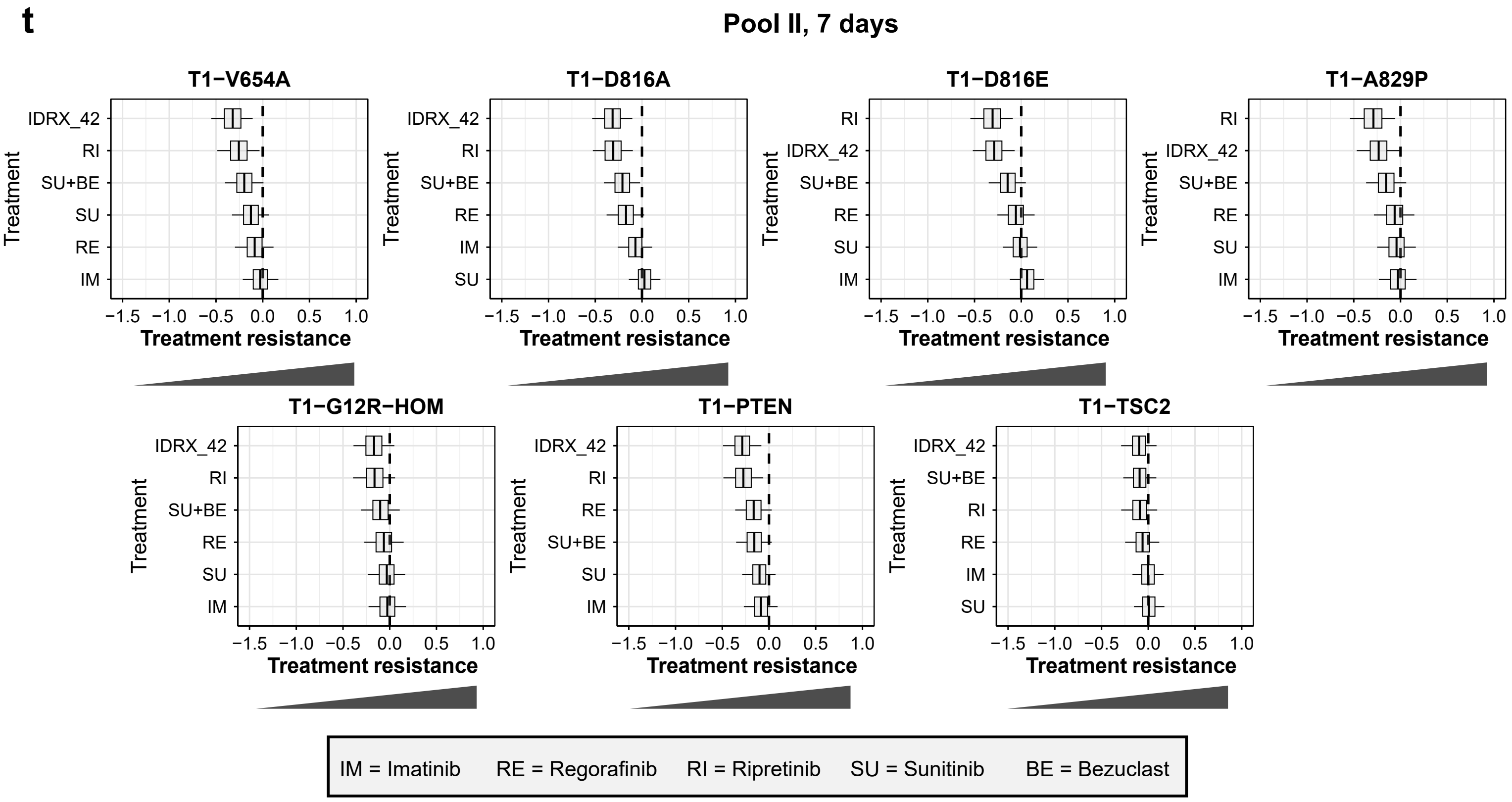

### Extended Data Fig. 5 - page 6/7

u

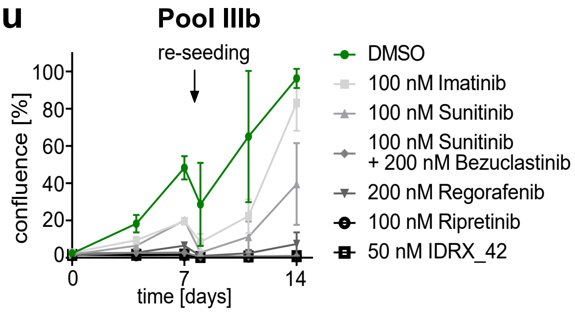


v

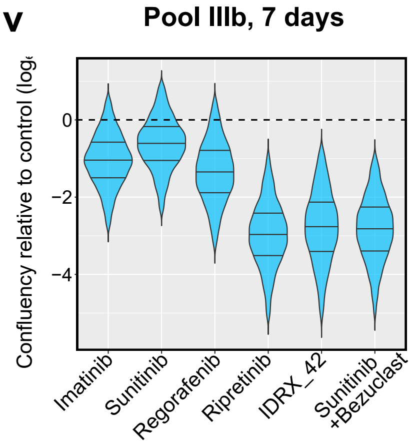


w

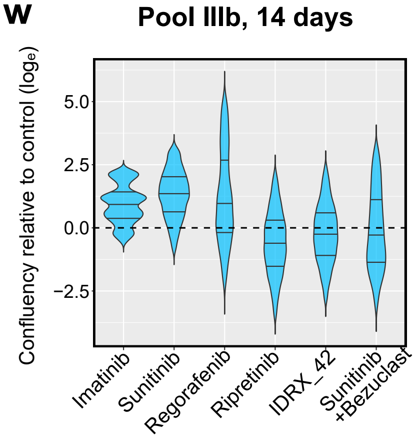


x

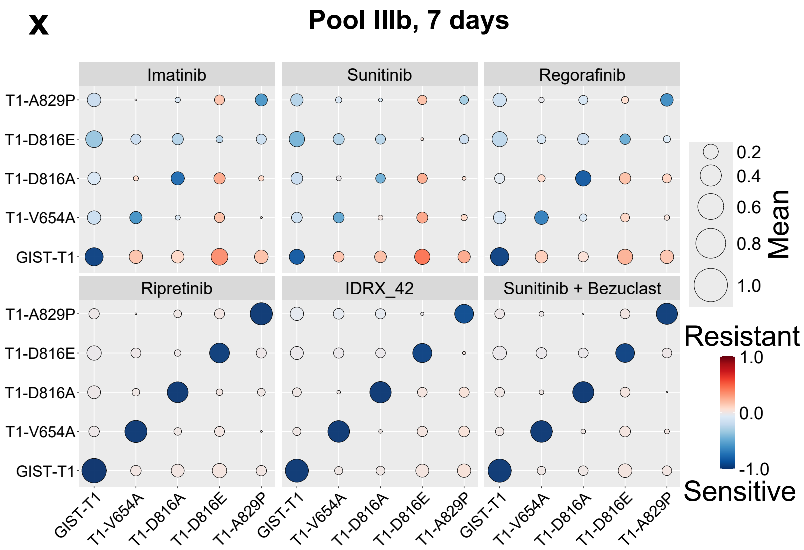


y

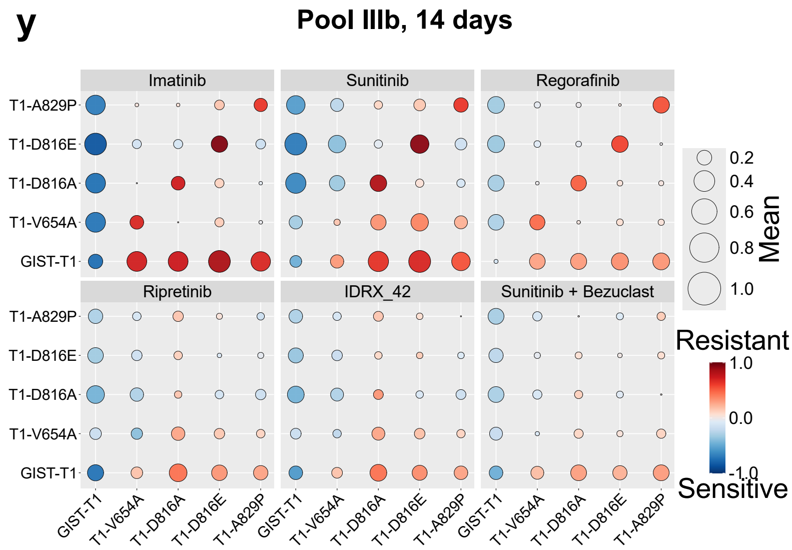

#### Extended Data Fig. 5 - page 7/7

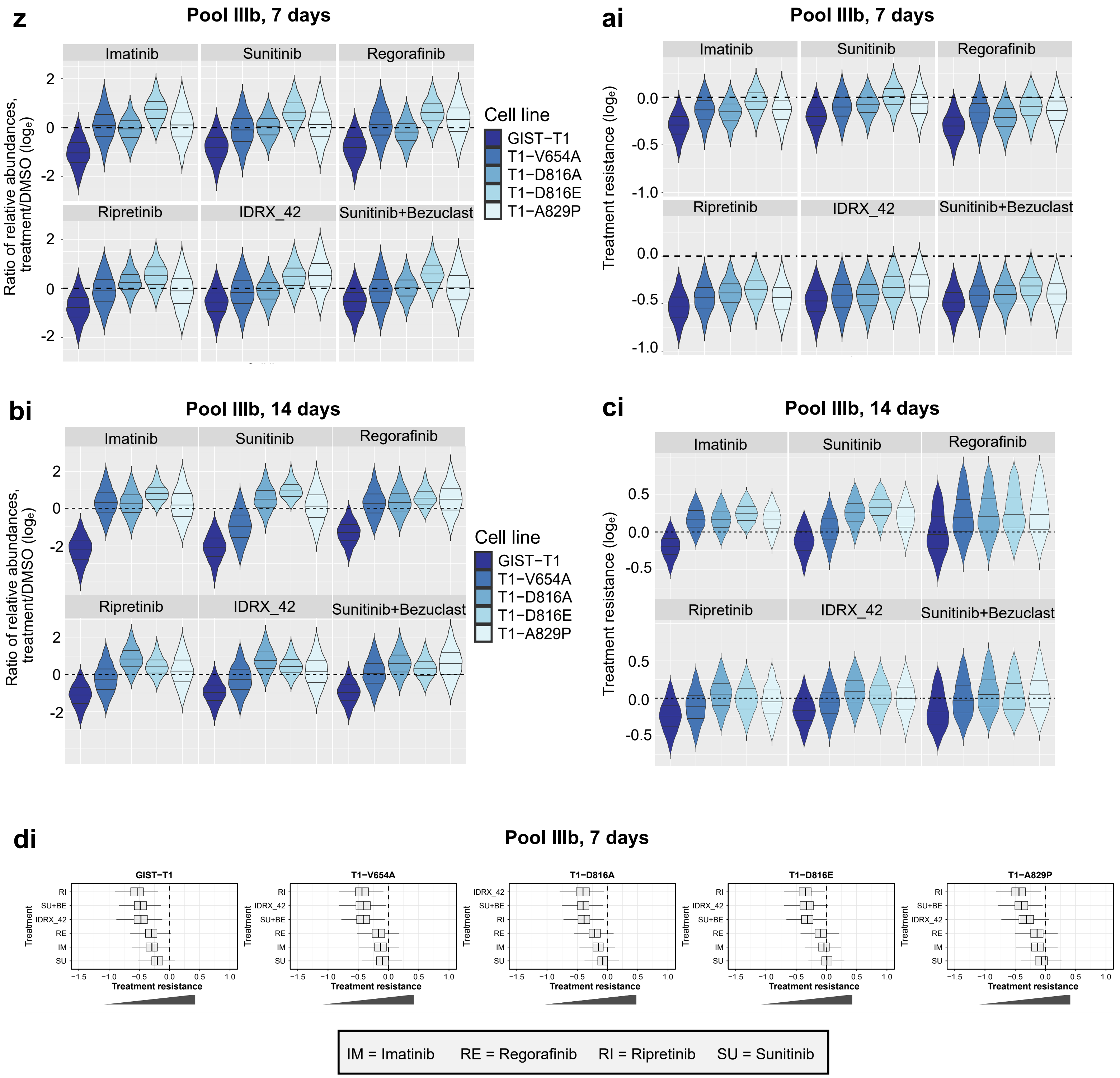

**Extended Data Fig. 5: Quantitative assessment of *in vitro* long-term treatment responses**

**a**) Confluencies of barcoded cell line in Pool I under treatment over time in triplicate wells – compare Main Fig. 5a. For better visibility, data was split over 4 individual graphs with repeated display of the DMSO control but was performed and analyzed in one combined experiment.

**b-e) Relative confluency responses (b, c) and genotype-specific pairwise comparisons of quantitative treatment resistance (d, e) in Pool I after 7 days (b, d) or 14 days (c, e) of *in vitro* treatments b, c)** Natural-log scaled confluency ratios treated to control, with uncertainty estimated using a Bayesian model. Horizontal black dashed line marks no change of treatment relative to control. Treatments along horizontal axis **d, e)** Bubble heatmaps (as in Extended Data Fig. 3a) are pairwise comparisons of treatment resistance among cell lines across individual treatment conditions in Pool I after 7 and 14 hours.

**f-i) Genotype-specific quantitative treatment responses in Pool I after 7 days (f, g) or 14 days (h, i) of *in vitro* treatment as indicated in b and c**

See caption Main Fig. 3d).

**j)** Ranked QTRs (as in Main Fig. 4) for 7 and 14 days of treatment.

**k-o) Confluencies (k, l, m) and genotype-specific pairwise comparisons of quantitative treatment resistance (n, o) in Pool II after 7 days (l, n) or 14 days (m, o) of *in vitro* treatment as indicated in k).**

**k)** Confluencies as explained in Main Fig. 5a). **l, m)** Confluency ratios as explained in Main Fig. 3b). **n, o)** Bubble heatmaps as explained in Extended Data Fig. 3a).

**p-s) Genotype-specific responses in Pool II after 7 days (p, r) or 14 days (q, s) of *in vitro* treatment as indicated in k).**

**p, q)** Natural-log scaled ratio of relative abundances as in Main Fig. 3c). **r, s)** Quantitative treatment resistance as in Main Fig. 3d).

**t)** Ranked QTRs as in Main Fig. 4.

**u)** See caption of Extended Data Fig. 3a).

**v-y) Relative confluency responses (v, w) and genotype-specific pairwise comparisons of treatment resistance (x, y) in Pool IIIb after 7 days (x, z) or 14 days (y, ai) of *in vitro* treatment as indicated in u).**

**v, w)** See caption of Main Fig. 3b). **x, y)** Bubble heatmaps as explained in Extended Data Fig. 3a).

**z-ci) Genotype-specific responses in Pool IIIb after 7 days (z, bi) or 14 days (ai, ci) of *in vitro* treatment as indicated in u).**

**z, bi)** As explained in Main Fig. 3c). **ai, ci)** As explained in Main Fig. 3d).

**di)** As explained in Main Fig. 4.

### Extended Data Fig. 6 - page 1/6

#### *in vivo* data for pool I

**a**

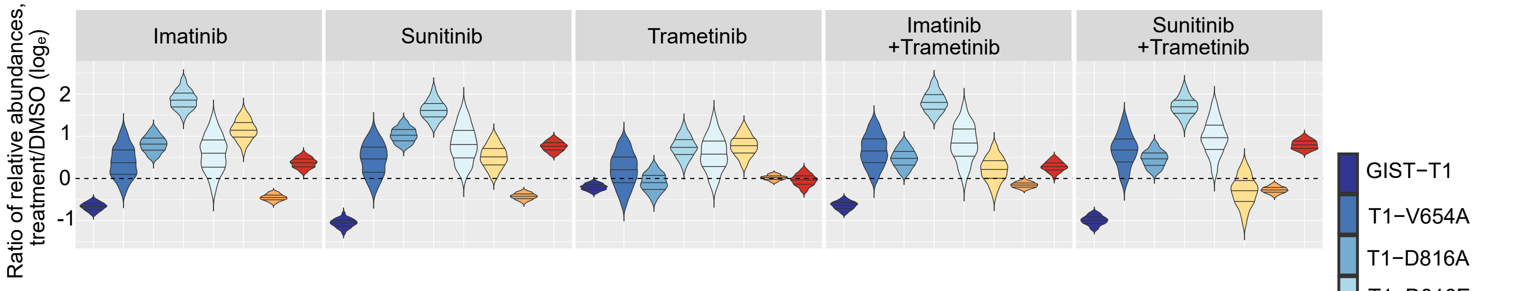


**b**

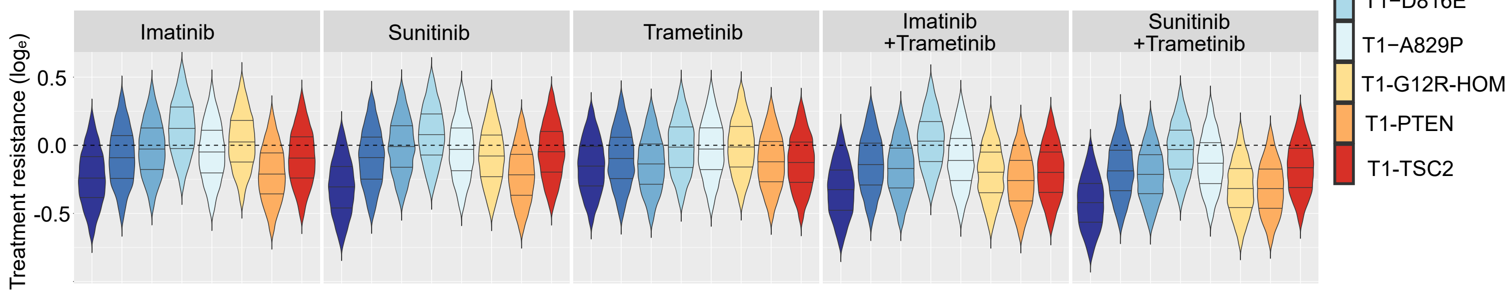


**c**

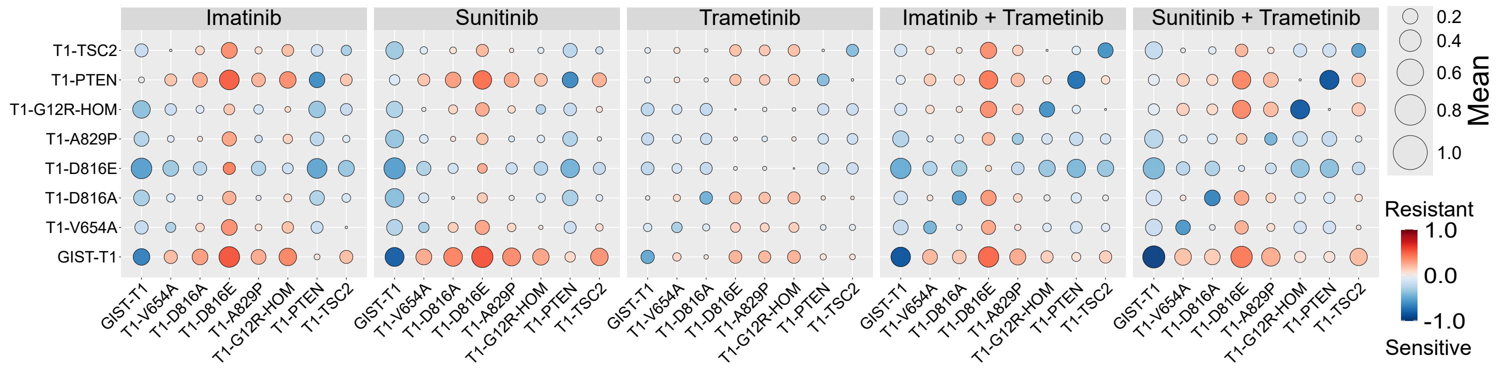


**d**

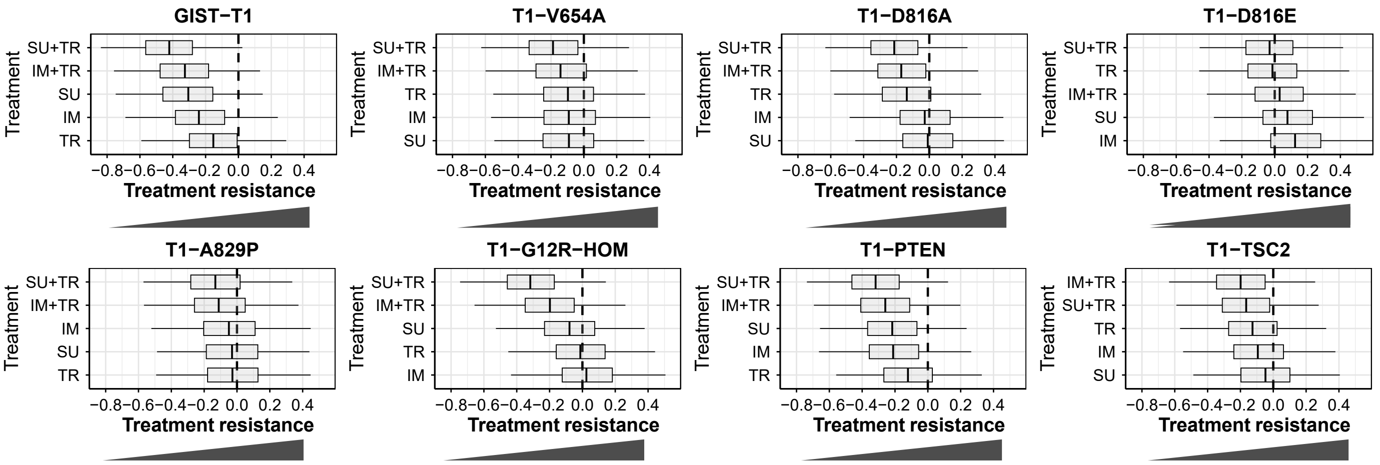


IM = Imatinib TR = Trametinib SU = Sunitinib

**Extended Data Fig. 6 - page 2/6**

***in vivo* data repetition of experiment for pool I**

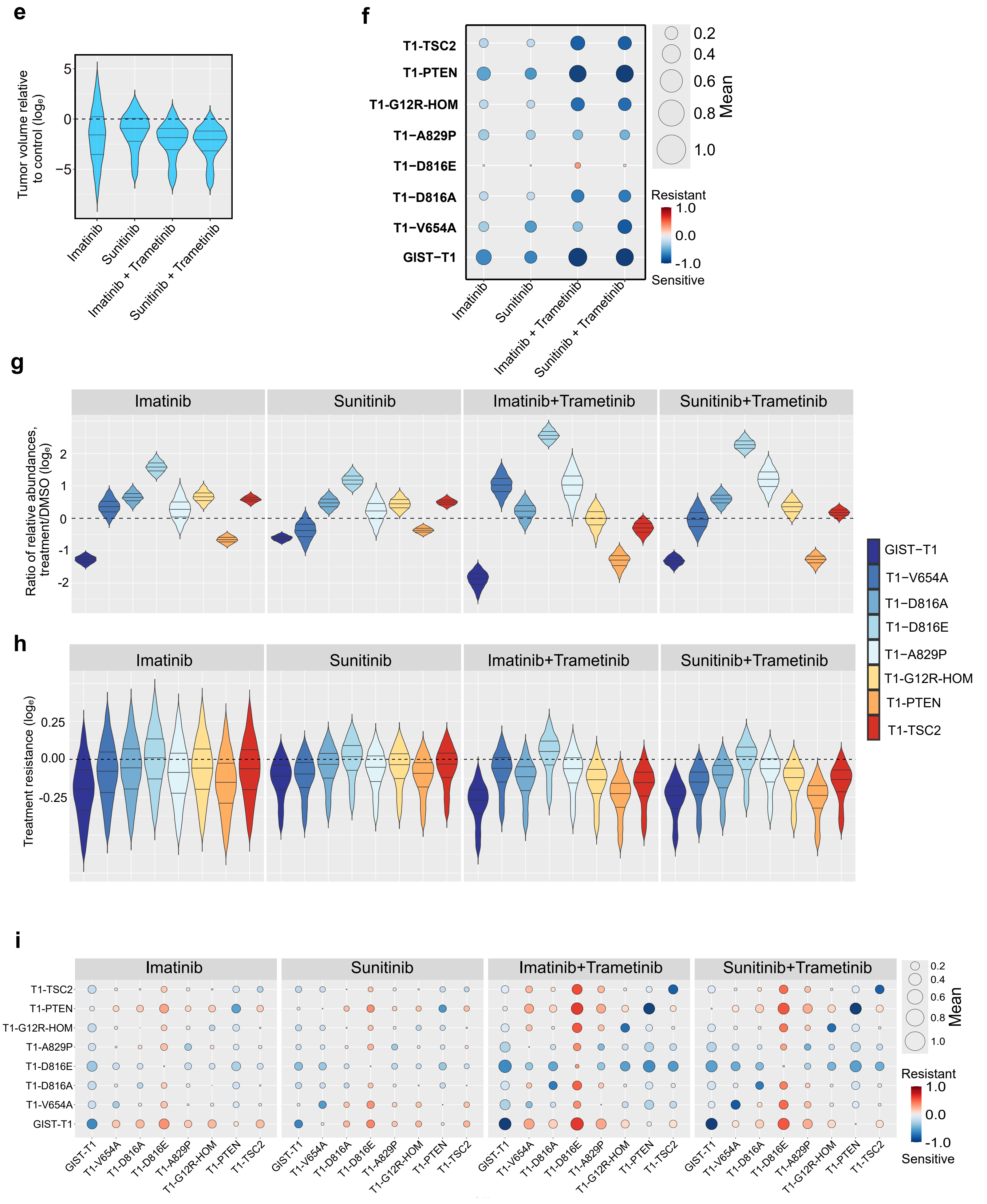

#### Extended Data Fig. 6 - page3/6

#### *in vivo* data repetition of experiment for pool I

**i**

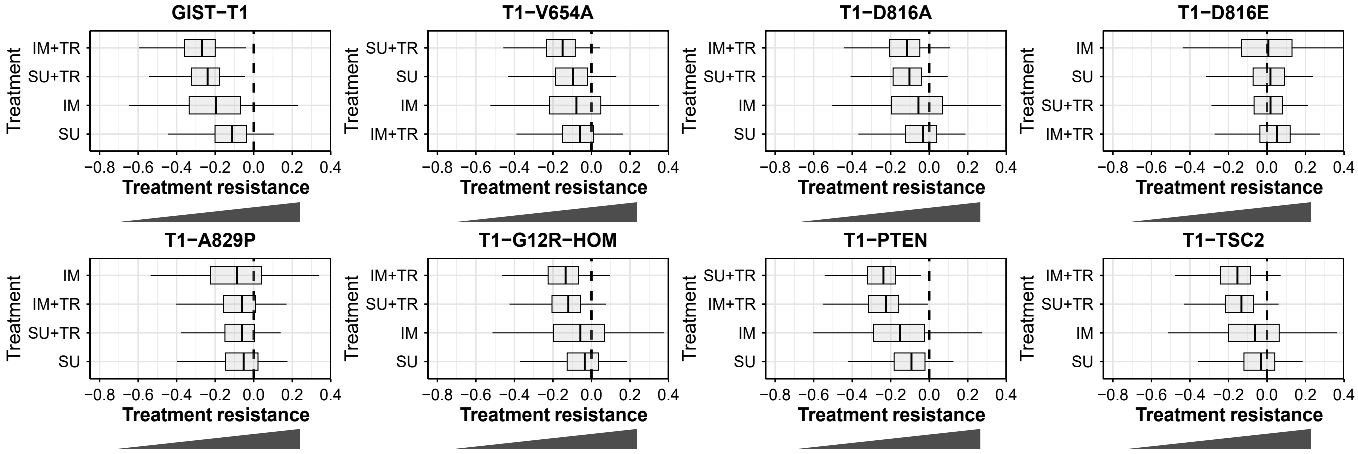


IM = Imatinib TR = Trametinib SU = Sunitinib

### Extended Data Fig. 6 - page 4/6

#### *in vivo* data for pool II

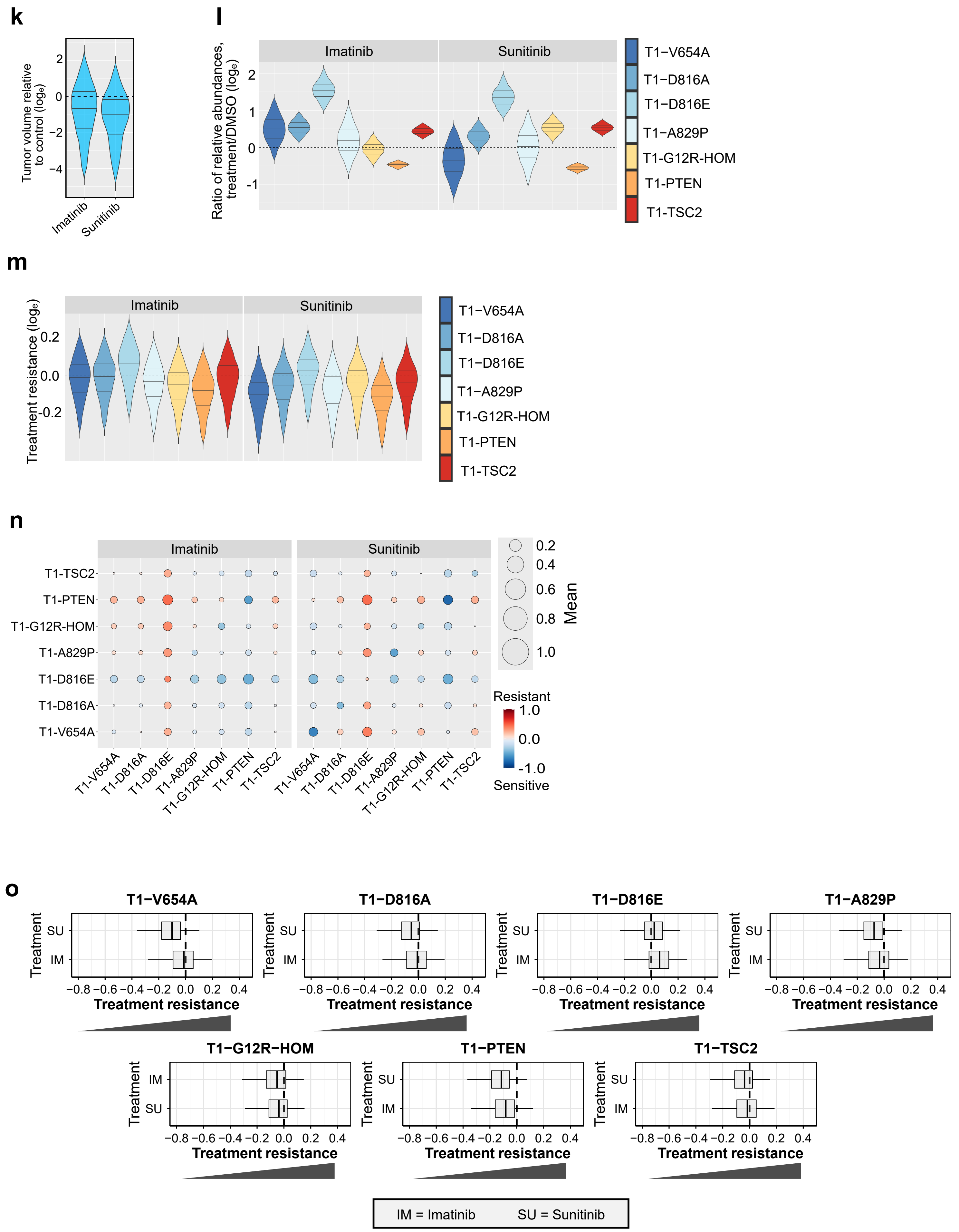

### Extended Data Fig. 6 - page 5/6

#### *in vivo* data for pool IIIb

**p**

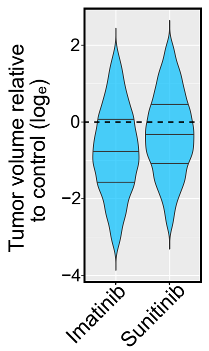


**q**

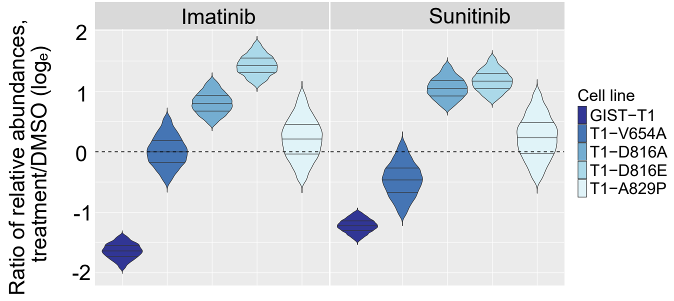


**r**

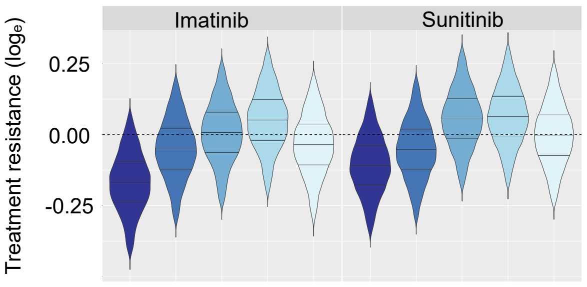


**s**

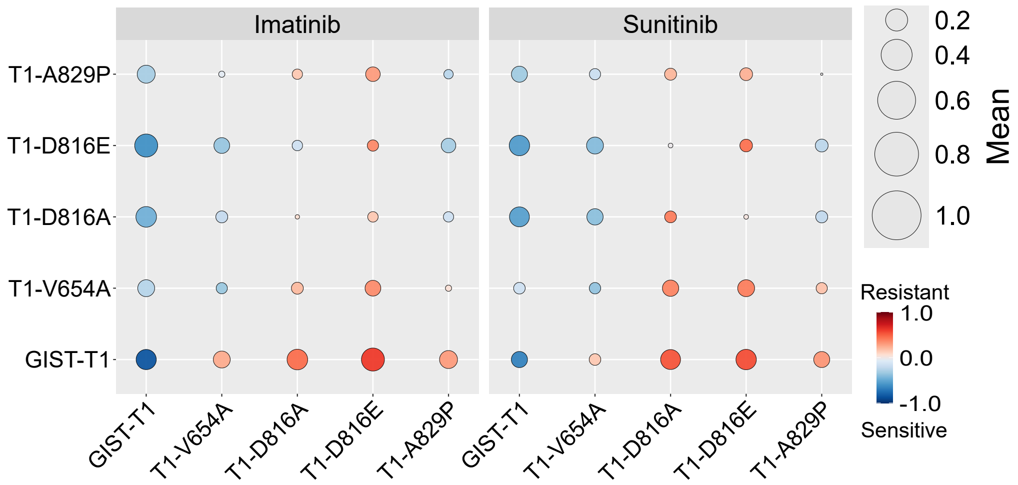


**t**

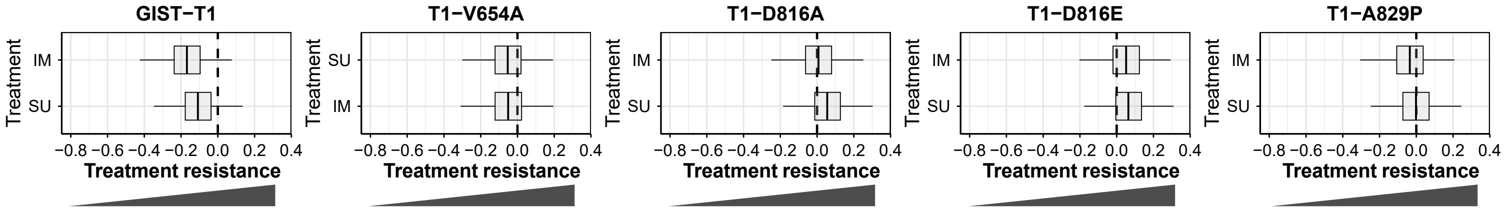


IM = Imatinib SU = Sunitinib

**Extended Data Fig. 6: BARMIX enables quantitative preclinical modelling *in vivo***

**a–d) Quantitative treatment responses *in vivo* for Pool I. Experimental setup and treatment concentrations are indicated in main Fig. 6a and b.**

**a)** As in Main Fig. 3c). **b)** As in Main Fig 3d). **c)** As in Extended Fig. 3a). **d)** As in Main Fig. 4.

**e–j) *In vivo* quantitative treatment responses for the independently repeated analysis of Pool I. Experimental setup and treatment concentrations are indicated in main Fig. 6a and b.**

**e)** Natural-log scaled ratio of tumor volume treated to control with uncertainty from Bayesian analysis. Treatments along bottom. **f)** Summary of QTR across genotypes and treatments as in Main Fig. 3e). **g)** As in Main Fig.c). **h)** QTRs from *in vivo* data as in Main Fig. 3d). **i)** Bubble heatmaps as in Extended Data Fig. 3a). **j)** Ranked QTRs as in Main Fig. 4.

**k–o) *In vivo* treatment responses for Pool II. Experimental setup and treatment concentrations are indicated in main Fig. 6a and b.**

**k)** As in Extended Data Fig. 6e). **l)** As in Main Fig. 3c). **m)** QTRs as in Main Fig. 3d). **n)** As in Extended Data Fig. 3a). **o)** As in Main Fig. 4.

**p–t) *In vivo* treatment responses for Pool IIIb. Experimental setup and treatment concentrations are indicated in main Fig. 6a and b.**

**p) As** in Extended Data Fig. 6e). **q)** As in Main Fig. 3c). **r)** QTRs as in Main Fig. 3d). **s)** Bubble heatmaps as Extended Data Fig. 3a). **t)** Ranked QTRs as in Main Fig. 4.
